# Cross-species and tissue imputation of species-level DNA methylation samples across mammalian species

**DOI:** 10.1101/2023.11.26.568769

**Authors:** Emily Maciejewski, Steve Horvath, Jason Ernst

## Abstract

DNA methylation data offers valuable insights into various aspects of mammalian biology. The recent introduction and large-scale application of the mammalian methylation array has significantly expanded the availability of such data across conserved sites in many mammalian species. In our study, we consider 13,245 samples profiled on this array encompassing 348 species and 59 tissues from 746 species-tissue combinations. While having some coverage of many different species and tissue types, this data captures only 3.6% of potential species-tissue combinations. To address this gap, we developed CMImpute (Cross-species Methylation Imputation), a method based on a Conditional Variational Autoencoder, to impute DNA methylation for non-profiled species-tissue combinations. In cross-validation, we demonstrate that CMImpute achieves a strong correlation with actual observed values, surpassing several baseline methods. Using CMImpute we imputed methylation data for 19,786 new species-tissue combinations. We believe that both CMImpute and our imputed data resource will be useful for DNA methylation analyses across a wide range of mammalian species.

## Introduction

DNA methylation is an epigenetic mark in which a methyl group is added to a cytosine. It is associated with gene regulation and disease^1,2^ and is a biomarker for individual characteristics such as age^3,4^. There is thus extensive interest in profiling DNA methylation in humans^5,6^ as well as other species^4,7–9^. In addition to studying DNA methylation profiles in individual species, insights have been gained from comparative epigenomic analyses across species as epigenetic information from one species will likely be informative to another species^10–16^. Along with varying at the species level, DNA methylation levels typically vary significantly across different tissue types and thus associate with cell and tissue identity.

Various methods exist for profiling DNA methylation data in biological samples, including microarrays^17–19^ and sequencing based assays such as whole genome bisulfite sequencing^20^ (WGBS) and reduced representation bisulfite sequencing^21^ (RRBS). Microarrays typically profile fewer cytosines than sequencing based assays, but allow for easier and more robust data collection and thus remain a popular approach to profile DNA methylation^22^. However, historically profiling DNA methylation using microarrays for species other than human or more recently mouse was limited due to the lack of applicable microarrays^18,23^. This recently changed with the development of the mammalian methylation array, which has array probes that allow the measurement of DNA methylation across mammalian species at a set of 36k CpGs that are well conserved across mammals^12^. This array has been used by the Mammalian Methylation Consortium to profile DNA methylation samples in at least one tissue type for over 348 mammalian species, collectively covering over 50 different tissue types^10,11^. However, the biological samples were gathered opportunistically and thus the collected data has an incomplete and imbalanced tissue type representation across species. For certain species, like horses and human, data from many tissue types were collected. However, for many other species, data from only one or two tissue types were collected. This results in experimental data being available for only a small percentage of the potential species-tissue combinations. The incomplete and imbalanced coverage of the experimental data thus motivates the need for computational approaches to accurately impute a DNA methylation sample representing a species and tissue type combination for which there is no experimental data available.

Current methods have shown that large-scale imputation of missing CpG sites and entire epigenetic datasets^24–30^ is effective in single-species datasets. However, since these existing methods are only designed to use data from a single species for imputation they are not able to leverage emerging large compendia of cross-species DNA methylation data^12,14^. In particular, in cases in which there is no data available in a given tissue type for a target species, methods that only consider data from a single-species would not be able to make predictions for that tissue type while methods that consider cross-species information could. Such an approach would have the potential to obtain predictions for human in an inaccessible tissue type by leveraging data from model organisms for that tissue type. A challenge towards large-scale cross-species DNA methylation imputation, besides having sufficient species and tissue coverage, has also been having sufficient coverage of a common set of CpGs across species. However, with the large compendium of methylation data for conserved CpGs that have been profiled with the mammalian methylation array, such data is now available.

To harness compendia of newly available cross-species methylation data to impute methylation values of shared CpGs across species for missing species and tissue combinations, we developed CMImpute (Cross-species Methylation Imputation). CMImpute specifically imputes samples representing a species’ mean methylation within a specific tissue type, henceforth referred to as a *species-tissue combination mean sample* or for short *combination mean sample*. Given the association of DNA methylation with species-level characteristics and cell and tissue identity, these types of combination mean samples have proven useful in cross-species epigenetic studies^10,11,16^. CMImpute takes as input exclusively methylation data with corresponding species and tissue labels to output combination mean samples. To perform species-tissue combination mean imputation, CMImpute uses a neural network architecture called a Conditional Variational Autoencoder (CVAE), an extension of the Variational Autoencoder (VAE). We note that VAEs and CVAEs have been used in various bioinformatics applications^31,32^ including in the context of DNA methylation^28,30^. Previous applications in the context of DNA methylation include imputing missing CpG values within an existing sample^28^ and generating additional human cancer DNA methylation samples when data for that cancer type is already experimentally available for some individuals^30^. However, none of these existing VAE and CVAE-based approaches have been designed for or applied in the context of cross-species DNA methylation imputation.

We demonstrate that CMImpute is able to accurately impute combination mean samples of missing species-tissue combinations through a cross-validation analysis of mammalian methylation array data. We show that imputed samples strongly correlate with observed species-tissue combination mean samples for held out combinations, when considering both samples across all probes and probes across all samples. In addition, we train CMImpute using all available observed samples from 746 species-tissue combinations to impute 19,786 mean samples representing the remaining 96.4% of combinations of the 348 species and 59 tissue types that had not been previously experimentally profiled. We demonstrate that the imputed samples, both from the cross-validation analysis and the full imputation, maintain inter-combination mean sample correlation patterns related to species and tissue types that are present in observed combination mean samples. Furthermore, we show how the imputed combination mean samples can be used to study the relationship between DNA methylation and maximum lifespan. The DNA methylation imputed by CMImpute vastly expands the coverage of species-tissue combination mean samples providing a resource for cross-species epigenetic studies or studies within a species lacking coverage of tissue types of interest.

## Results

### Overview of CMImpute

CMImpute takes as input individual methylation samples, spanning a common set of CpGs, and the corresponding species and tissue label for each sample. We note there can be multiple training samples representing the same species and tissue combination since samples from more than one individual are collected for most species-tissue combinations that have observed data available. CMImpute outputs imputed species-tissue combination mean samples for combinations with no observed samples available but where other tissues were profiled in the target-species and other species were profiled in the target-tissue (Fig. 1a, Supplementary Data 1). To capture inter- and intra-species tissue signals for imputation, CMImpute trains a neural network using the input methylation samples and species and tissue labels. Using the trained neural network, CMImpute then imputes the methylation level for each CpG in missing species-tissue combinations (Fig. 1b).

**Figure 1.**
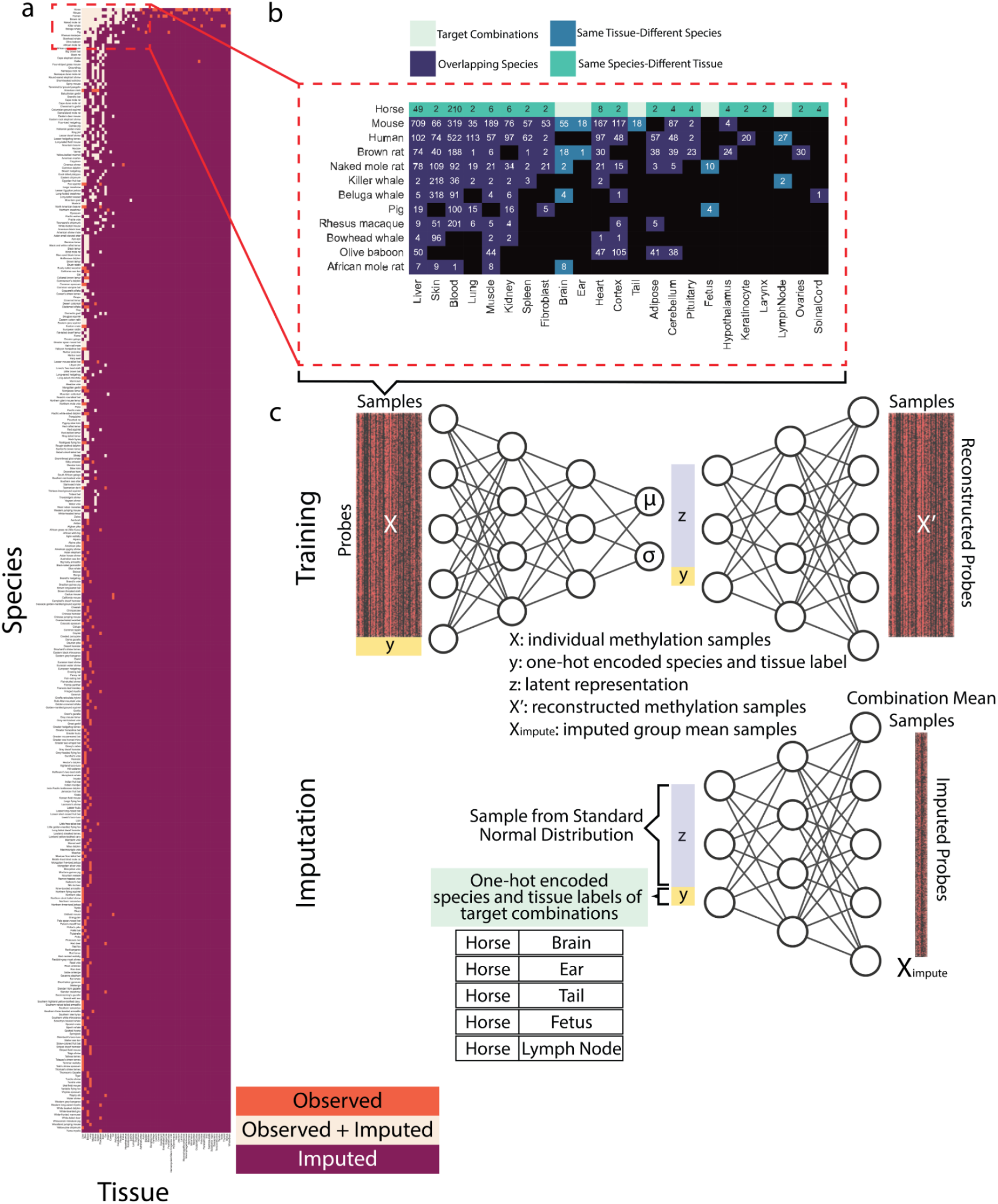
Data and method overview. **a)** Grid of all species-tissue combinations colored by what type of data is now available for each combination (observed, imputed or both). Observed combinations were observed in at least one individual and have no imputed data available. Combinations with both observed and imputed data available (Observed+Imputed) were both observed in at least one individual and had predictions generated for it in cross-validation. Imputed combinations represent all combinations without observed data and were included in the final imputed data compendium only. Species are sorted top-to-bottom by the number of available tissues. Tissues are sorted left-to-right by the number of available species. Species and tissues available in the same number of tissues or species, respectively, listed in alphabetical order. Number of individual samples in each species and tissue type (listed in same order as figure) available in Supplementary Data 1. Subset of species and tissues outlined in the dotted red line highlighted for use in (b). **b)** Example displaying the different categories of data used during training by CMImpute. In this example, there are no observed samples for certain horse tissues (target combinations). Samples from three categories of training data are used as input for CMImpute: target species data from non-target tissues (same species-different tissue), data from other species in the target tissues (same tissue-different species), and data from overlapping tissues between the target species and other species if available (overlapping species). For this example, the target species is horse and the target tissues are brain, ear, tail, fetus, and lymph node. **c)** Method overview illustrating the neural network architecture used for *Training* and *Imputation.* CMImpute’s CVAE framework takes as input a matrix of individual observed samples with corresponding species and tissue labels. During *Training*, the CVAE learns methylation patterns from the three categories of training data. Once trained, *Imputation* can occur. The CVAE uses the learned parameters to impute species-tissue combination mean samples of the missing target species-tissue combinations. In the example illustrated, CMImpute imputes the missing horse tissues.

The specific neural network CMImpute uses to perform the imputation is a CVAE, an extension of the VAE. A VAE is a self-supervised neural network architecture trained to reconstruct the original input and regularized to maintain a probabilistic latent space^33^. This regularization enables VAEs to both encode an input sample into and to generate a new sample from its latent space^33^. However, VAEs do not have control over the types of data generated. CVAEs extend the VAE framework by adding labels corresponding to each input sample^34^. These labels provide additional information about each sample during training and allow for control over the generated samples.

### CMImpute predictions qualitatively agree with observed data

To assess CMImpute’s imputation performance we first applied it to a subset of the species and tissues for which there was mammalian methylation array data available. Specifically, we applied CMImpute in 5-fold cross-validation to impute data for 465 combination mean samples for which we also have observed data available. These 465 combination mean samples correspond to 134 species with data from more than one tissue type available and 23 tissues with data from more than one species available. We compared CMImpute’s performance to the performance of four baseline methods (Methods). One baseline was logistic regression, where for each probe we applied logistic regression with the species and tissue labels as the features. Another baseline was a global baseline which was the mean of all training samples. The other two baselines were species and tissue baselines, which were based on the mean of training samples within the same species or the same tissue, respectively.

We first qualitatively evaluated CMImpute’s predictions by generating heatmaps that show the methylation values for each combination mean sample and probe after applying hierarchical clustering with optimal leaf ordering^35^. We did this both for all probes (Fig. 2a-c, Supplementary Fig. 1a) and a subset of 11,749 probes that are mappable to a unique genomic location in most mammalian species, referred to as highest-coverage probes (Methods) (Supplementary Fig. 1b). The samples mainly clustered by phylogenetic order with tissue clustering primarily occurring within the orders. We compared these heatmaps to corresponding heatmaps based on observed data. The CMImpute and species baseline-imputed heatmaps appeared similar to the observed methylation patterns at the inter-species levels. However, when the species contribution was removed from the observed and imputed datasets by subtracting the average of all same-species training samples (Fig. 2d-f, Supplementary Fig. 1c), we observed differentially methylated regions in the observed and CMImpute combination mean samples but not the species baseline-imputed samples. This lack of tissue signal in the species baseline was expected as it was defined as the average of all available same-species samples. The logistic regression and tissue baseline, while appearing to capture tissue-specific methylation patterns, did not appear to effectively capture the observed species-specific methylation patterns (Supplementary Fig. 1a,c).

**Figure 2.**
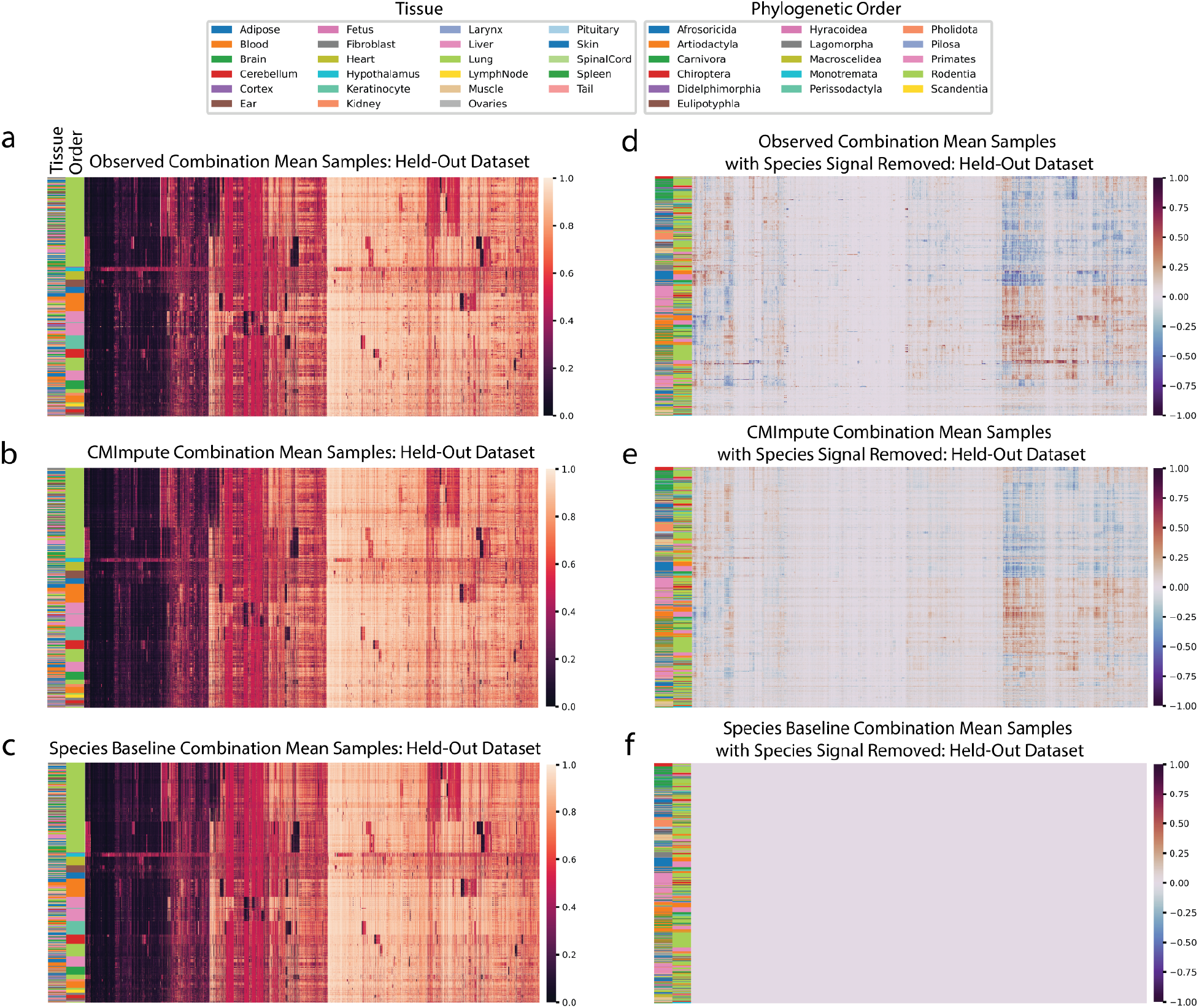
Visualization of imputed species-tissue combination mean samples relative to held-out observed values. a-c) Heatmaps of methylation probe values for the **a)** observed data held-out during cross-validation and **b)** CMImpute’s and **c)** the species baseline’s predictions of the held-out data (additional baselines are shown in Supplementary Fig. 1a). Each row is a species-tissue combination mean sample and each column is a methylation probe. Samples and probes were ordered based on hierarchical clustering followed by optimal leaf ordering. Color bars on the left indicate the phylogenetic order (inner) and tissue (outer) corresponding to the samples. Legends corresponding to the color bars are below the heatmaps. Color scale representing methylation values from 0 to 1 on the right. **d-f)** Heatmaps of the **d)** observed, **e)** CMImpute, and **f)** species baseline datasets with the species signal removed to highlight the differentially methylated tissue regions (observed tissue AUC score of 0.850, CMImpute tissue AUC score of 0.874). Species signal was removed by subtracting the average methylation values of same-species training samples from the full methylation values displayed in a. Color scale representing methylation delta values from -1 to 1 on the right.

### Analysis of combination mean sample-wise imputation performance

We next quantitatively evaluated the imputation performance of CMImpute predictions generated in 5-fold cross-validation relative to the baselines. For this, we evaluated the agreement of CMImpute and baseline-imputed species-tissue combination mean samples with the corresponding held-out combination mean samples using the average Pearson correlation and mean squared error (MSE). On average when considering all probes, CMImpute combination mean samples had a 0.920 correlation, compared to 0.906 for the species baseline, 0.886 for logistic regression, 0.778 for the tissue baseline, and 0.803 for the global baseline (Fig. 3a). To further put the agreement of CMImpute’s predictions with held-out data in context, we also computed average pairwise correlations between samples of the same species and tissue combination, which was 0.981 (Fig. 3a). This suggests that there still exists some reproducible biological signal not captured by CMImpute’s predictions. When considering the subset of highest-coverage probes, CMImpute’s performance increased to 0.932 and continued to be greater than the species baseline’s performance of 0.897, logistic regression’s performance of 0.923, tissue baseline’s performance of 0.880, and global baseline’s performance of 0.877 (Supplementary Fig. 2a). Similar performance trends were also seen using MSE as the evaluation metric (Supplementary Fig. 2b-c).

**Figure 3.**
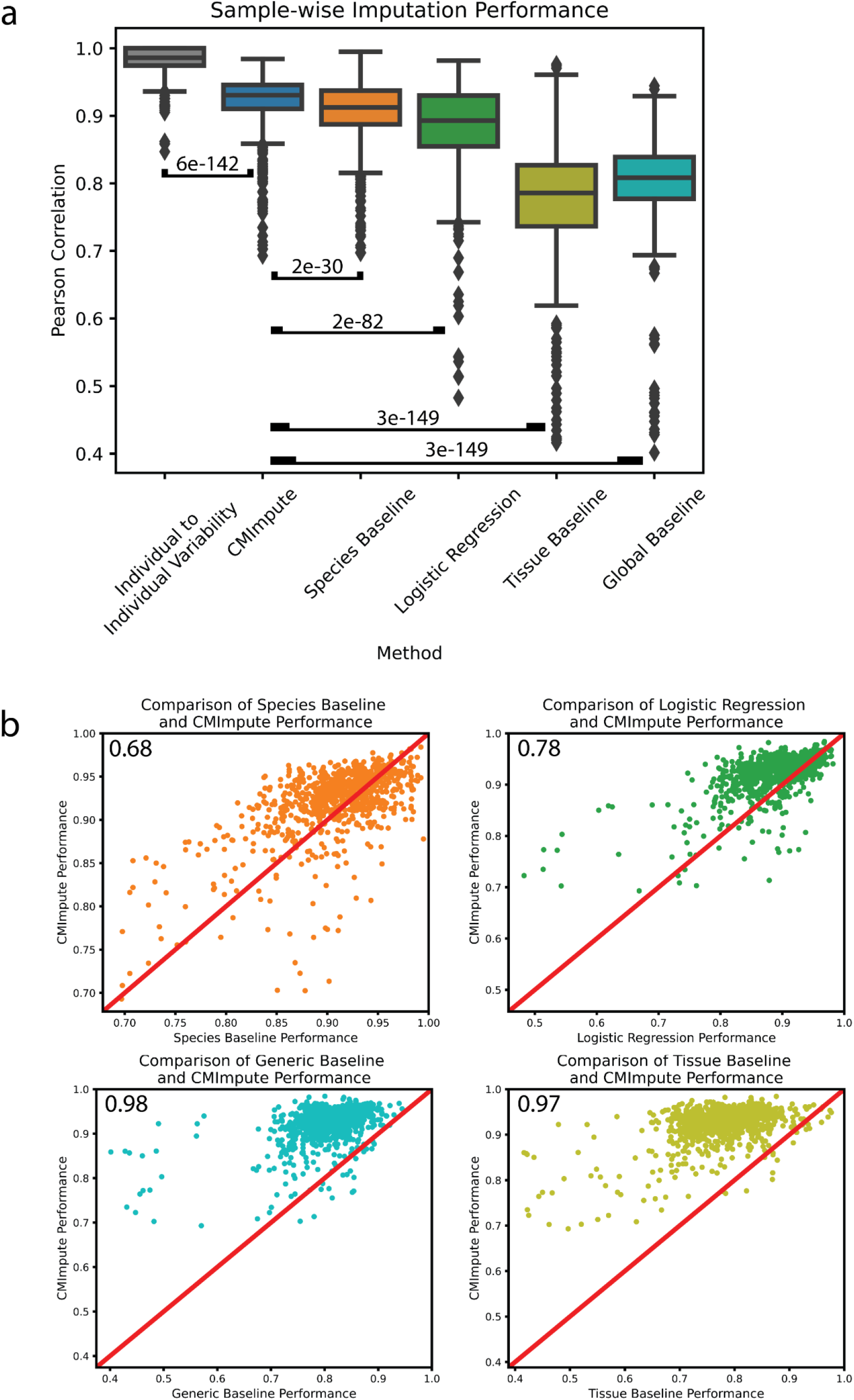
Sample-wise performance of imputed species-tissue combination mean samples. **a)** Sample-wise Pearson correlation of imputed species-tissue combination mean samples with held-out observed values when considering all methylation probes. Individual to individual variability represents the average pairwise correlation of observed data between individuals of the same species and tissue type for each combination. Baselines and individual to individual variability labeled by Wilcoxon signed-rank test p-value comparing CMImpute’s sample-wise Pearson correlation to the individual to individual variability and each baseline’s sample-wise Pearson correlation for each imputed species-tissue combination ([CMImpute, Individual to Individual Variability], [CMImpute, Species Baseline], [CMImpute, Logistic Regression], [CMImpute, Tissue Baseline], [CMImpute, Overall Baseline]). **b)** Comparison of CMImpute and baseline imputation performance measured via sample-wise Pearson correlation with held-out observed data across all probes. The y-axis is CMImpute’s performance on each imputed combination. The x-axis is the species baseline’s (top left), logistic regression’s (top right), tissue baseline’s (bottom left), or overall baseline (bottom right) performance on each imputed combination. Each dot is a single imputed species-tissue combination mean sample. The red diagonal line represents equal performance between CMImpute and the baseline. If a point is above the diagonal, CMImpute outperforms the baseline on the corresponding imputed combination mean sample and vice versa. Values in the upper left corners are the fractions of samples where CMImpute outperforms the corresponding baseline.

In addition to having higher mean correlation with the observed data than baselines, CMImpute also had higher correlation for a large majority of individual species-tissue combination mean samples (Fig. 3b, Supplementary Data 2,3). Specifically, CMImpute outperformed the species baseline in 68% of species-tissue combination mean samples, in 78% compared to logistic regression, in 98% compared to tissue baseline, and in 97% compared to the global baseline based on sample-wise Pearson correlation across all probes. Combination mean samples where the species baseline or logistic regression had higher correlation than CMImpute were for combinations for which there was overall a relatively low number of individual samples from the target species represented in the training data (Supplementary Fig. 3a-b). CMImpute additionally outperformed the baselines for the large majority of samples when restricting to the subset of highest-coverage probes and when considering MSE instead of correlation (Supplementary Table 1). These results demonstrate that CMImpute is able to impute species-tissue combination mean samples for held-out combinations with greater accuracy than the baselines for a large majority of species-tissue combinations.

### Analysis of probe-wise imputation performance

In addition to computing sample-wise performance, which was based on the agreement of probe values within the same combination mean sample, we also quantified probe-wise performance, which was based on agreement of probe values across samples. For this, we again conducted evaluations in 5-fold cross-validation using both the Pearson correlation coefficient and MSE. We note that for probes that have almost no variance across combination mean samples, we expect the Pearson correlation to be less informative.

We first focused on the subset of highest-coverage probes since within this subset variation in probe methylation values across samples would less likely be driven by differences in mappability across species. For this subset, CMImpute had a mean probe-wise correlation of 0.623 significantly outperforming the species, logistic regression, tissue, and global baselines of 0.518, 0.494, 0.217, and 0.002, respectively (Fig. 4a). When considering all probes, CMImpute’s mean probe-wise correlation of 0.688 was also higher compared to the species, logistic regression, tissue, and global baselines of 0.650, 0.545, 0.142, and 0.004 respectively (Fig. 4b). We saw similar performance trends using mean MSE for both all probes and the subset of highest-coverage probes (Supplementary Fig. 4). We note that when considering the median as opposed to the mean correlation and all probes the species baseline did have a higher median correlation of 0.716 compared to CMImpute’s 0.703. However, this was not the case for other baselines or for the highest-coverage probes where the median correlations for the species baseline and CMImpute were 0.535 and 0.626, respectively (Fig. 4a,b).

**Figure 4.**
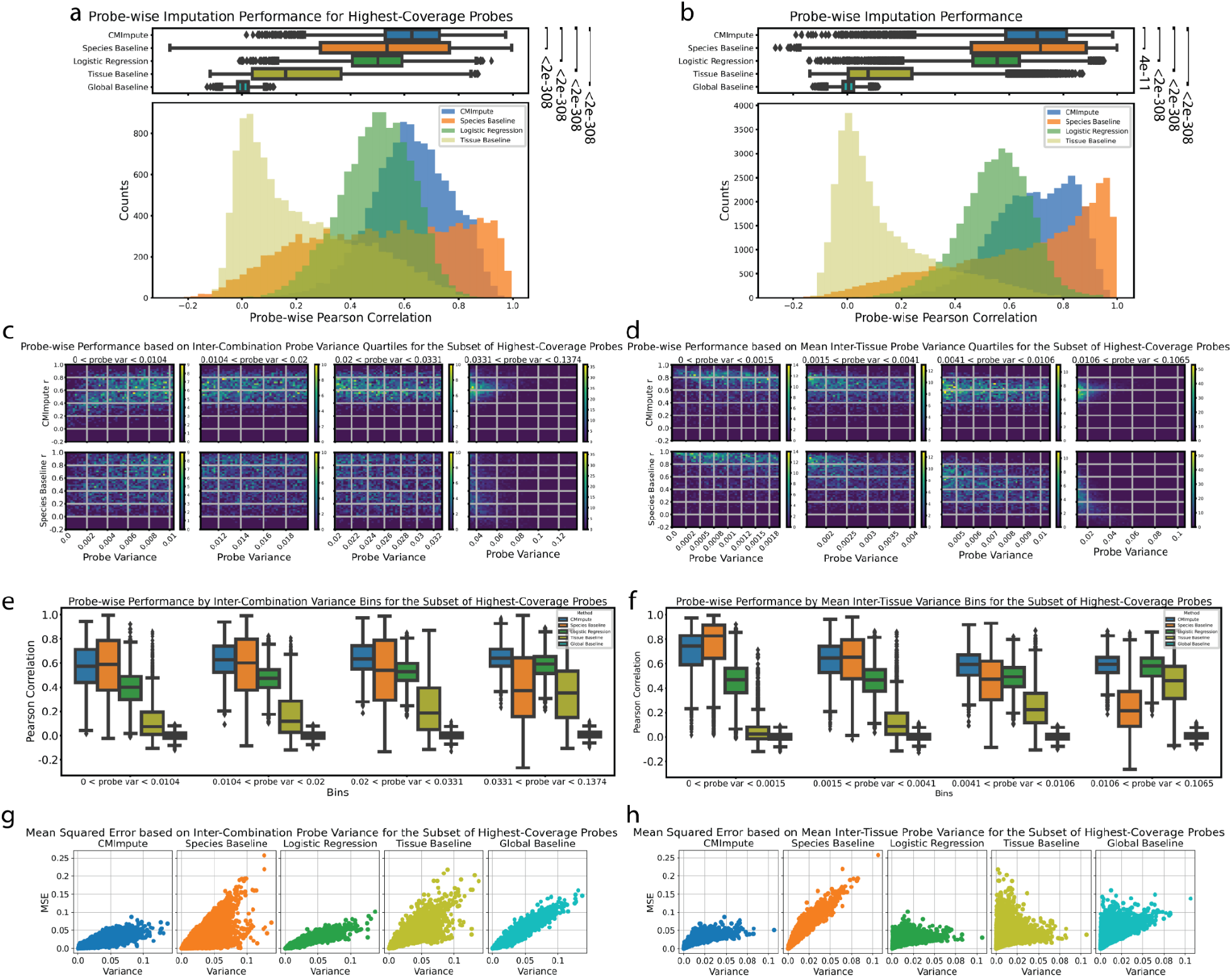
Probe-wise performance of imputed species-tissue combination mean samples. **a)** Distributions of probe-wise Pearson correlations with held-out observed values when considering highest-coverage probes. The top boxplots show the distribution of probe-wise correlations with held-out observed values. The bottom histograms show the number of imputed combination mean samples across 50 performance bins. As the global baseline predictions do not vary within a fold, the probe-wise performance is not meaningful and this is not included in the histograms. Legend for both boxplots and histograms shown in histogram plot. Baselines labeled by Wilcoxon signed-rank test p-value comparing CMImpute’s probe-wise Pearson correlation and each baseline’s probe-wise Pearson correlation for each imputed probe ([CMImpute, Species Baseline], [CMImpute, Logistic Regression], [CMImpute, Tissue Baseline], [CMImpute, Global Baseline]). Corresponding plots for subsets of higher variance probes can be found in Supplementary Fig. 9a-c. **b)** Same as a) except for all probes. Corresponding plots for subsets of higher variance probes can be found in Supplementary Fig. 9d-f. **c)** 2-d histograms showing CMImpute (top row) and species baseline (bottom row) probe-wise correlation as a function of inter-combination probe variance when considering the subset of highest-coverage probes. Each row contains four heatmaps corresponding to variance quartiles. Within each variance quartile, the heatmap shows the number of probes with a certain probe variance along the x-axis and certain probe-wise correlation along the y-axis split into 50 bins along each axis. Each quartile is labeled with its own color bar. Color bar scales for each quartile are consistent across methods. Corresponding plots when considering all probes can be found in Supplementary Fig. 5e. Remaining baselines can be found in Supplementary Fig. 6d. **d)** Same as c except for mean inter-tissue variance when considering the subset of highest-coverage probes. Corresponding plots when considering all probes can be found in Supplementary Fig. 6a. Remaining baselines can be found in Supplementary Fig. 6c. **e-f)** Boxplots of the probe-wise Pearson correlation with held-out observed values for the subset of highest-coverage probes in each **e)** inter-combination and **f)** mean inter-tissue variance quartile. Each variance quartile represented in the boxplots correspond to the variance quartile in the 2-d histograms from c-d. c-f demonstrate that CMImpute yields relatively consistent performance across variance quartiles and outperforms the baselines, particularly on higher variance probes. Corresponding plots when considering all probes can be found in Supplementary Fig. 5f,6b. **g)** Probe-wise MSE (y-axis) relationship to inter-combination variance (x-axis) when considering the subset of highest-coverage probes shown in separate plots for CMImpute, species baseline, logistic regression, tissue baseline, and overall baseline (left to right). The x-axis is the probe variance. Each dot corresponds to a single imputed probe. **h)** Same as g except for mean inter-tissue variance. Mean inter-species variance can be found in Supplementary Fig. 10a. Corresponding plots for all probes can be found in Supplementary Fig. 10b. CMImpute yields relatively low MSE across lower and higher variance probe levels and outperforms the baselines, particularly on higher variance probes.

We further analyzed probe-wise performance as a function of probe variance, allowing us to compare imputation performance across differentially and non-differentially methylated regions. We used three types of variances: inter-combination variance, mean inter-tissue variance, and mean inter-species variance (Methods). Inter-combination variance represented the mean probe variation between different species-tissue combinations. Mean inter-tissue variance represented the average probe variation between tissues within a species. Mean inter-species variance represented the average probe variation between species within a tissue.

We first analyzed the probe-wise Pearson correlations of CMImpute-imputed combination mean samples as well as those from the baselines with held-out data as a function of each type of variance. This revealed that for inter-combination and mean inter-species variance, CMImpute consistently had higher mean correlation across all variance quartiles. For mean inter-tissue variance, CMImpute consistently had a higher mean correlation in all but the lowest variance quartile where the correlation is expected to be less informative (Fig. 4c-f, Supplementary Fig. 5a-b,9a-c). Similar results were also seen when expanding to consider all probes with CMImpute outperforming the species baseline for most variance quartiles (Supplementary Fig. 5c-d,6a-b,7a-b,9d-f). When comparing CMImpute to the other baselines, it consistently had higher correlations across all quartiles based on the highest-coverage probes and all probes (Fig. 4e-f, Supplementary Fig. 5-9).

We additionally used probe-wise MSE to evaluate the imputation performance of CMImpute and the baselines as a function of variance. We first considered the subset of highest coverage probes (Fig. 4g-h, Supplementary Fig. 10a) and found that similar to the probe-wise Pearson correlation analysis, CMImpute had a lower mean MSE than the species baseline in all but the lowest mean inter-tissue variance quartile (Supplementary Fig. 11a,12a). Thus, while the species baseline relative to CMImpute more accurately imputes probes with low mean inter-tissue variance thus containing limited tissue-specific activity, it less accurately imputes probes of higher mean inter-tissue variance thus containing greater tissue-specific activity signal. CMImpute additionally had a lower mean MSE for all inter-combination and mean inter-species variance quartiles for the highest-coverage probes (Supplementary Fig. 11b-c,12b-c). Similar results as with the highest-coverage probes were also seen in the highest and second highest variance quartiles when expanding to consider all probes, with CMImpute consistently outperforming the species baseline in these quartiles (Supplementary Fig. 10b,11d-f,12d-f).

When comparing CMImpute to logistic regression, CMImpute consistently had lower MSEs with an average margin of 0.0158 across all quartiles and variance types when considering all probes. When considering the subset of highest-coverage probes, CMImpute had lower MSEs in the first three mean inter-tissue quartiles with an average margin of 0.00127 across all quartiles. When comparing CMImpute to the tissue and global baselines, CMImpute consistently had lower MSEs across all quartiles when considering all probes and the highest-coverage probes (Fig. 4g-h, Supplementary Fig. 10-12). Overall these results demonstrate that CMImpute has effective performance according to probe-wise imputation metrics.

### Impact of amount of available data on imputation accuracy

We also sought to understand how the amount of available training data impacted imputation performance. We first investigated the sample-wise performance as a function of the number of tissue types within a target species. We note this evaluation does not consider the amount of data available in the tissue types for non-target species. Consistent with results based on all imputed combination mean samples, when investigating subsets of imputed samples based on the amount of available training data CMImpute also outperformed the baselines in sample-wise imputation performance evaluations using mean Pearson correlation (Supplementary Fig. 13). As the number of tissue types in the target species increased, CMImpute’s mean imputation performance with held-out data tended to increase (*r*= 0.181), with the performance increasing from 0.915 mean correlation for one tissue type in the target species to 0.951 for five tissue types. We did not observe a corresponding increase for the mean correlation between individual observed samples within the same tissue and species combination (r= -0.048), which had values of 0.982 and 0.978 for one and five tissue types in the target species, respectively (Fig. 5a). This result is consistent with CMImpute’s improved performance with additional tissue types being driven by the additional available training data and not differences in the observed variability across individuals within the species and tissue combinations.

**Figure 5.**
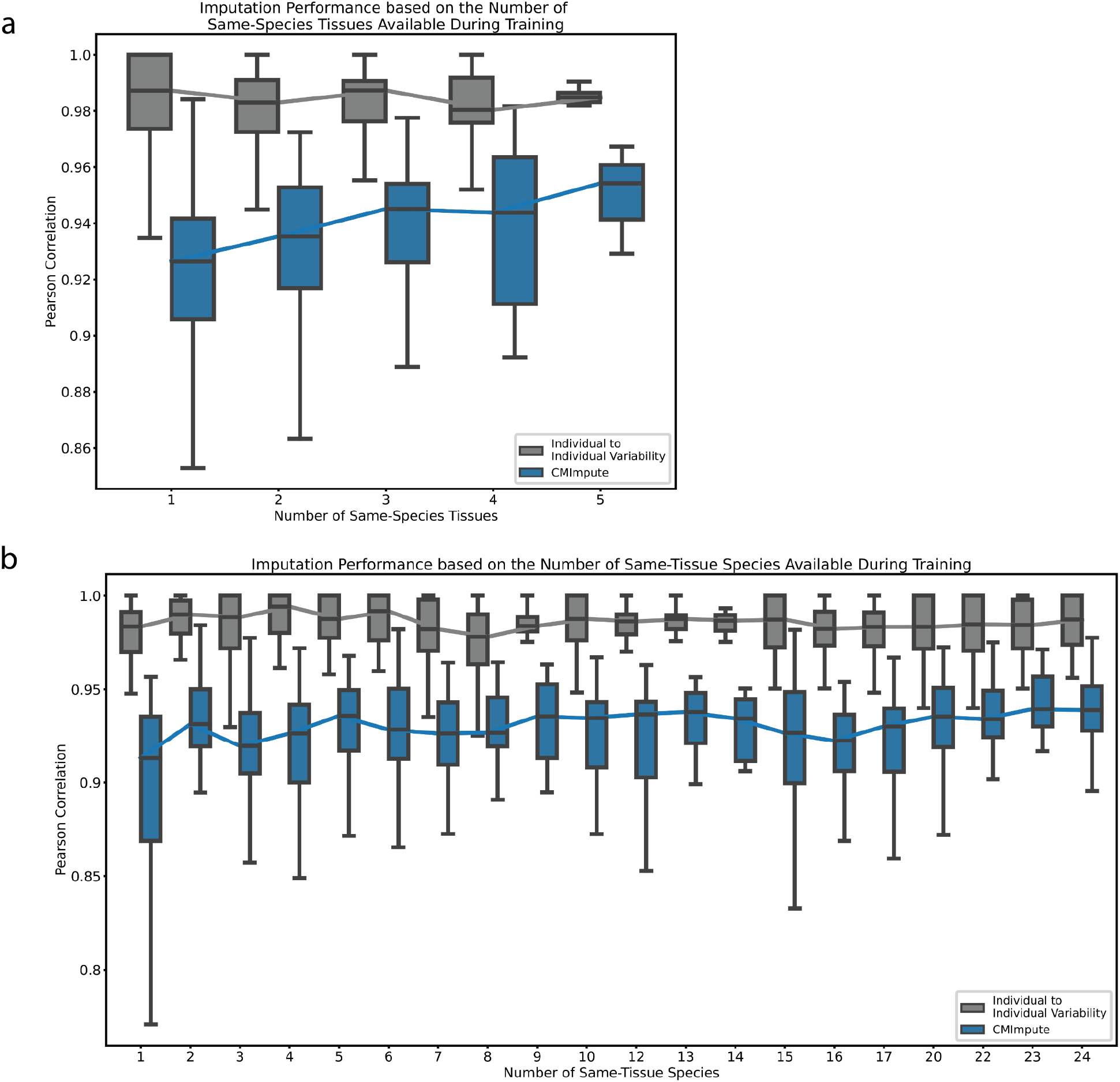
Impact of the amount of available training data on CMImpute performance. **a)** Sample-wise Pearson correlation distributions as a function of the number of tissue types available in the target species during training. The box plot shows the distribution of Pearson correlation for each number of tissue types. Line connects the median correlations for an imputation method across all tissue type counts. Individual to individual variability represents the average pairwise correlation of observed data between individuals of the same species and tissue type for each combination. Baseline performance can be found in Supplementary Fig. 13a. **b)** Similar to a, but sample-wise Pearson correlation as a function of the number of species available during training in the target tissue. Baseline performance can be found in Supplementary Fig. 13b.

We also evaluated the sample-wise performance as a function of the number of different species within the target tissue. We note this evaluation does not consider the number of available tissue types in the target species. Consistent with the results from the evaluation as a function of the number of tissue types in the target species, CMImpute outperformed the baselines in sample-wise imputation performance evaluations as a function of the number of species for the target tissue type (Fig. 5b). CMImpute’s performance increased as the number of same-tissue species increased (r=0.166), with the mean performance increasing from 0.893 to 0.932 when going from one to two same-tissue species and achieved the maximum performance of 0.938 when considering the maximum number of same-tissue species. Overall, CMImpute yielded high performance with limited amounts of same-tissue and same-species training data, but performance still increased with additional training data.

### Imputation of non-observed species and tissue combination mean samples

Using all the data collected using the mammalian methylation array that we are considering here (Supplementary Data 1), we applied CMImpute to impute all combinations not present in this input compendium (Methods). This resulted in imputed data for 19,786 species-tissue combination mean samples without observed data available spanning all 348 species and 59 tissue types (Fig. 1a imputed).

We first clustered and visualized heatmaps of the methylation values for all probes in the CMImpute species-tissue combination mean samples (Fig. 6a). Similar to what we previously observed when clustering based on the observed data (Fig. 2a), these heatmaps also showed sample clusters that corresponded to phylogenetic order. Also consistent with these phylogenetic order associated clusters, heatmaps of pairwise correlations between samples showed a correlated block structure between phylogenetic orders in both observed (Fig. 1a observed, Fig. 6b) and CMImpute-imputed (Fig. 1a imputed and observed+imputed, Fig. 6c) combination mean samples. We confirmed that these patterns could not be explained based on mappability differences between species as we saw similar patterns when we clustered and visualized the data restricted to the highest-coverage probes (Supplementary Fig. 14). For comparison, we also conducted a similar set of clustering and visualizations for the data imputed from the baseline methods (Supplementary Fig. 15-17). This showed that the logistic regression and tissue baselines did not show clear clustering of samples corresponding to species (Supplementary Fig. 15-17b-c), while as expected the species baseline did (Supplementary Fig. 15-17a).

**Figure 6.**
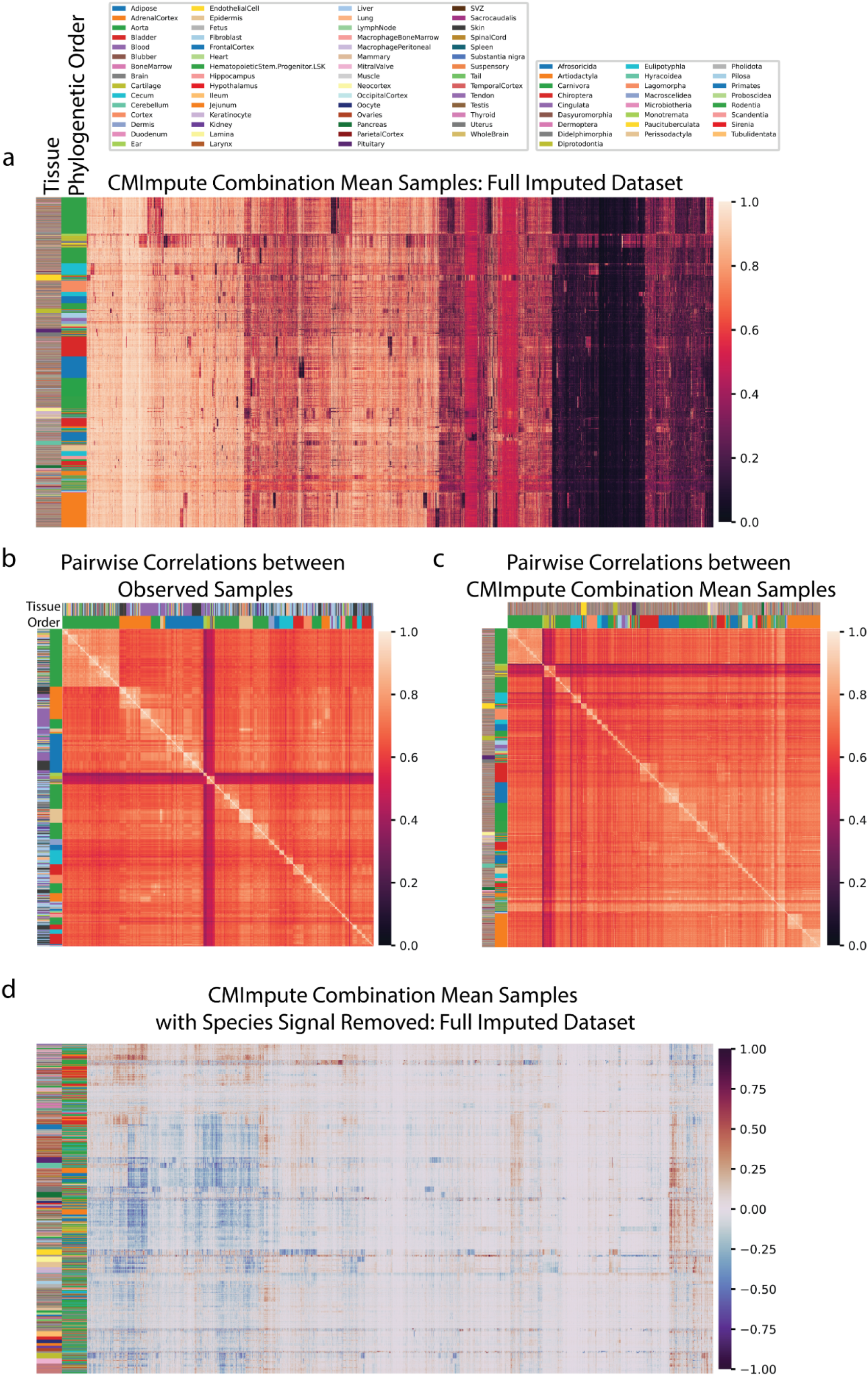
Visualization of CMImpute-imputed samples of non-observed combinations. **a)** Heatmap of the imputed dataset’s methylation probe values. Samples and probes were ordered based on hierarchical clustering followed by optimal leaf ordering. Color bars on the left indicate the phylogenetic order (inner) and tissue (outer) corresponding to the samples. Legends corresponding to the color bars can be found above the heatmaps. Color scale representing methylation values from 0 to 1 on the right. CMImpute-imputed combination mean samples of missing species-tissue combinations mainly cluster by phylogenetic order. **b-c)** Heatmaps of pairwise correlations between species-tissue combination mean samples for **b)** all 746 observed species-tissue combinations and **c)** 20,251 CMImpute-imputed samples from both the cross-validation analysis and full imputed compendium. Samples are ordered based on hierarchical clustering followed by optimal leaf ordering of the methylation samples (same order as a). Color bars on the left indicate the phylogenetic order (inner) and tissue (outer) corresponding to the samples. Despite the observed heatmap considering a small subset of the imputed datasets, both the observed and imputed heatmaps demonstrate highly correlated block structures between and within phylogenetic orders. **d)** Heatmaps of imputed dataset considered in a) with the species signal removed to highlight the differentially methylated tissue regions.

To highlight tissue-specific signal captured in CMImpute-imputed values, we also clustered and visualized the methylation values and pairwise correlations based on all probes after regressing out the species contribution (Fig. 6d). This revealed clusters of samples corresponding to the same or similar tissue types and a correlated block structure corresponding to tissue types (Supplementary Fig. 18a). For comparison we also conducted a similar analysis for the baseline methods (Supplementary Fig. 18b-e,19a-d). Unlike CMImpute, the species baseline did not capture tissue-specific methylation patterns (Supplementary Fig. 18b,19a). The tissue and logistic regression baselines, which previously did not show species-specific signal, did show tissue-specific methylation patterns (Supplementary Fig. 18c-d,19b-c). We note that many of those patterns were not consistent with patterns seen in either the observed data or CMImpute’s predictions.

### Quantifying species and tissue signals in combination mean samples

In addition to identifying species and tissue signals through clustering and visualization, we also directly quantified species and tissue signals in combination mean samples. We did this for species signal by evaluating the ability of pairwise correlations between species-tissue combination mean samples to predict whether a combination mean sample pair is of the same species quantified using an Area Under receiver operating characteristic Curve (AUC), and similarly for tissue signal based on whether the pair is of the same tissue, first using all probes (Fig. 7). We performed these evaluations on observed combination mean samples as well as imputed combination mean samples from CMImpute and the baseline methods. To directly compare the AUC values based on observed and imputed combination mean samples, we restricted this analysis to the species and tissue combinations included in the cross-validation analysis (Fig. 1a observed+imputed). The observed and CMImpute combination mean samples had similar tissue signals with AUC values of 0.656 and 0.667, respectively, and similar species signals with AUC values of 0.992 and 0.979, respectively. The tissue and species AUC values for combination mean samples based on logistic regression (0.750, 0.786) were higher and lower, respectively, than observed and CMImpute AUC values. As expected, the species baseline had a high species AUC value (0.993) and a low tissue AUC value (0.503), while the tissue baseline had a high tissue AUC value (0.857) and low species AUC value (0.471). To confirm that the species and tissue signals were not simply reflecting mappability differences between species, we additionally restricted this analysis to the subset of highest-coverage probes and saw similar trends (Supplementary Fig. 20a). In addition, we confirmed that when using all imputed combination mean samples (20,251 combinations considered in Fig. 6c), including those for which we did not have observed data, we saw similar tissue and species AUC values for CMImpute and the baselines (Supplementary Fig. 20b,c).

**Figure 7.**
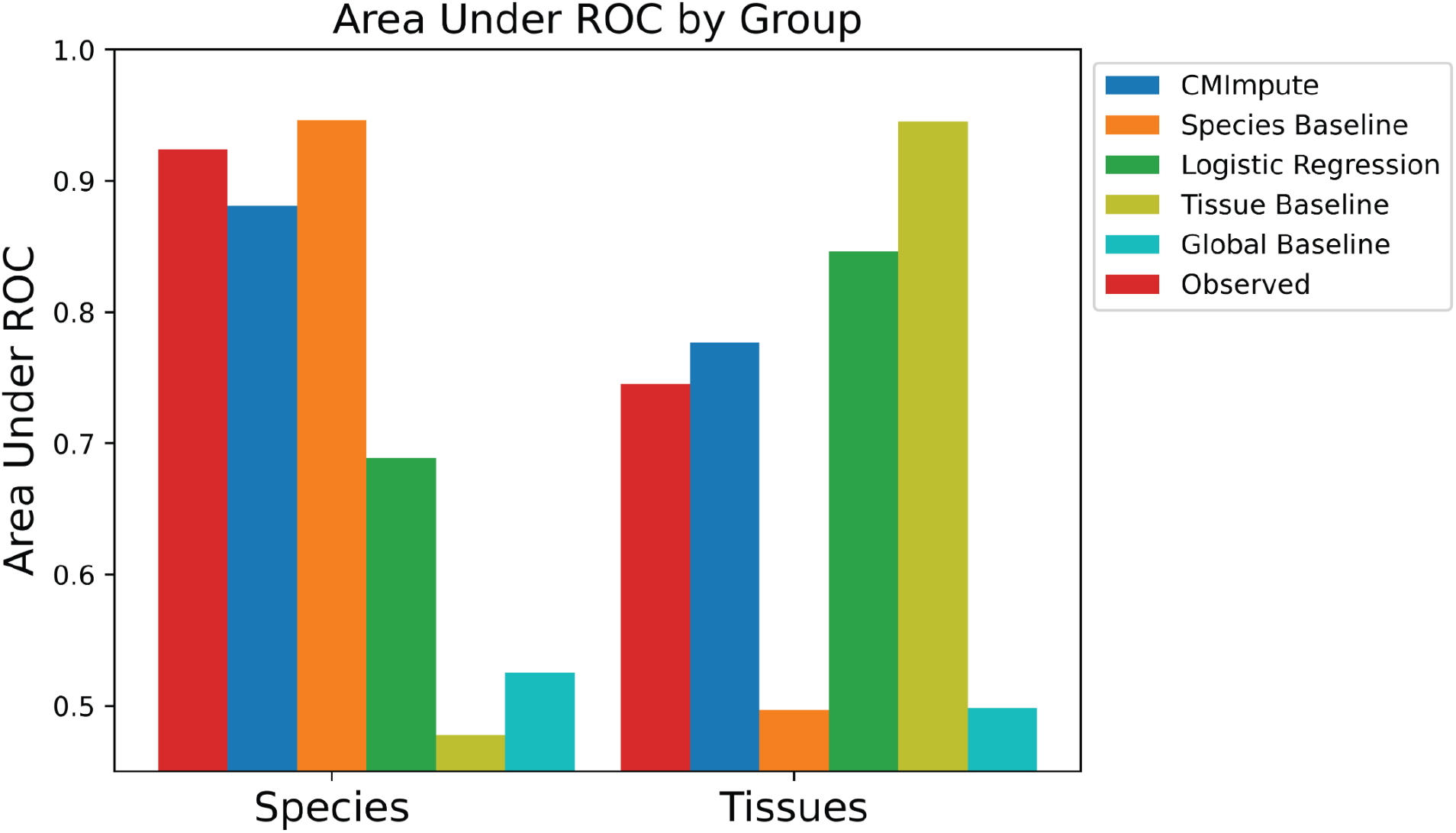
Species and tissue signal in observed and imputed samples. Area Under ROC values for predicting whether samples within the cross-validation dataset are from the same species or tissue based on their pairwise correlations for all probes. Corresponding plot for the subset of highest-coverage probes can be found in Supplementary Fig. 20a.

### Imputed species-tissue combination mean samples are predictive of a species’ maximum lifespan

We next applied the imputed species-tissue combination mean samples to analyze the relationship between species-level methylation values and species’ maximum lifespan^10,16^. For this we followed a similar approach to Li et al.^16^ and performed linear regression to predict the logarithm of a species’ maximum lifespan based on methylation data. Specifically, we first performed a linear regression analysis in a tissue-agnostic setting based on the average of combination mean samples within a species to see if similar predictive performance could be achieved with CMImpute’s imputed data for tissue types without observed data as could be with observed data for tissue types in which observed data was available (Methods).

We evaluated the predictive performance using Pearson correlation with log maximum lifespan using a leave-one-species-out (LOSO) analysis and saw similar correlations of 0.813 and 0.829 for the observed and imputed data, respectively (Fig. 8a,b). The MSE distributions were also similar with low median MSEs for both observed and imputed data of 0.064 and 0.047, respectively (Supplementary Fig. 21a). In addition to the imputed and observed data leading to similar predictive performance, the actual predicted values of the logarithm of maximum lifespan were also highly correlated with each other (0.973, Fig. 8c), demonstrating that CMImpute samples capture similar signals related to species maximum lifespan as the observed data.

**Figure 8.**
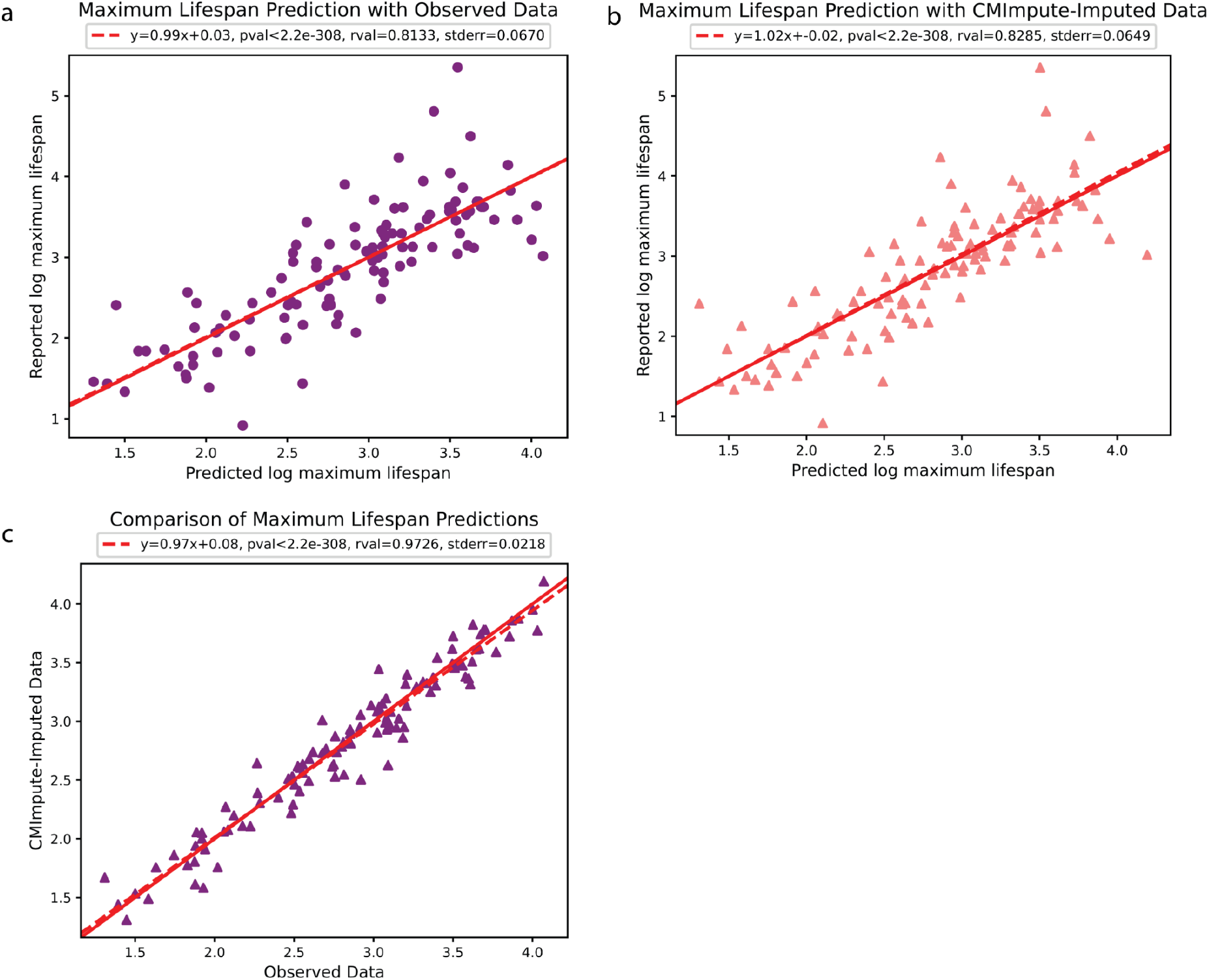
Prediction of species’ maximum lifespan using combination mean samples. a-b) Leave-one-species-out (LOSO) linear regression analysis using species-average samples to predict a species’ maximum lifespan. For each plot, each dot corresponds to a species. Dashed red line is the regression line between predicted and reported log-maximum lifespan. Solid red line denotes y = x. Regression coefficients, Pearson correlation, p-value, and standard error are shown above each plot. Average methylation calculated over **a)** exclusively observed methylation samples or **b)** CMImpute-imputed species-tissue combination mean samples. Predicted log-maximum lifespan (x-axis) plotted against the reported log-maximum lifespan (y-axis). **c)** Comparison of maximum lifespan predictions based on average species methylation samples between using observed and imputed data.

Similar results were also seen in a tissue-specific setting when considering individual tissue types (average Pearson correlation of 0.772 and 0.762 and median MSE of 0.072 and 0.064, for observed and imputed respectively, restricted to tissue types with observed data in at least three species, Supplementary Fig. 21b) (Methods).

## Discussion

Following the development and large-scale application of the mammalian methylation array^10–12^, there has been a large increase in available methylation data from a wide range of mammalian species. However, while high, though still incomplete, tissue coverage is present in certain species such as horse, mouse, and human, most species have limited profiled tissue types. Handling this incomplete and imbalanced tissue sample coverage across the 348 species in the Mammalian Methylation Consortium compendium^10–12^ presents a significant bioinformatics challenge. To tackle this, we introduced CMImpute, designed to estimate mean methylation values for various species-tissue combinations. We trained CMImpute on the data from the Mammalian Methylation Consortium^10–12^. CMImpute was specifically designed for imputing these species and tissue combinations that have not been previously experimentally profiled but where other tissues have been profiled in the target species and other species have been profiled in the target tissue types (Fig. 1b). To do this CMImpute uses a CVAE which has the advantages of being able to share information across probes and learn non-linear relationships.

Through a cross-validation analysis, we demonstrated that CMImpute accurately imputed combination mean samples of missing species and tissue combinations, outperforming multiple baselines both in terms of agreement with held-out observed data for sample and probe-wise performance. CMImpute still yielded reasonable performance that was better than the baselines when limited same-species and same-tissue information for the target species-tissue combination was available, though as expected performance increased with additional same-species or same-tissue information.

Finally, we trained CMImpute on all data from the mammalian methylation array that we were considering and used the subsequent model to impute 19,786 new species-tissue combination mean samples representing 348 species and 59 tissue types. We showed based on these predictions and the cross-validation predictions that CMImpute’s imputed samples contained species and tissue signals that were consistent with observed patterns. We also demonstrated that using the new imputed combination samples we could predict the maximum lifespan of a species with similar accuracy as when using observed samples.

While CMImpute already showed effective performance, there are possible extensions that could be investigated in future work. Currently CMImpute does not account for other sample attributes besides species and tissue, however a potential avenue for future work is to investigate if extending CMImpute with additional one hot-encoded labels corresponding to additional attributes such age, sex, or individual donor performs effectively. CMImpute also does not currently explicitly account for phylogenetic information, which could potentially be used to improve predictive performance. CMImpute makes its predictions based on exclusively methylation data as opposed to incorporating sequence or other biochemical data so as not to confound downstream analyses with other layers of information. Furthermore, in the context of highly conserved sites across species, limited species-specific variation could be predicted from sequence, and other sources of biochemical data in the target species are often not available.

However, future work could investigate approaches for also incorporating sequence information or other biochemical data, when available, into predictions. Finally, this work was limited to applying CMImpute to data from the mammalian array, while there is also now a large-scale cross-species methylation dataset based on RRBS^14^. However we note such data, unlike the mammalian array, does not specifically target highly conserved regions across mammals. Future work could also investigate applying and possibly extending CMImpute to RRBS or other methylation assays.

Even with these avenues for future work, CMImpute already has effective performance for imputing species-tissue combination mean samples, which enabled us to provide computational predictions that vastly expand the current compendium of methylation information. We expect CMImpute and its imputed datasets will be a resource for comparative epigenetic studies analyzing species and tissue-level methylation patterns across mammalian species.

## Methods

### Mammalian methylation array data

We used a dataset of 13,245 individual DNA methylation samples across 348 mammalian species and 59 tissues spanning 746 unique tissue-species combinations^10,11^ (Supplementary Data 1). All of this data was generated by the Mammalian Methylation Consortium on the developed mammalian methylation array^22^ and corresponded to the subset of the consortium data available at the time of our analyses. The array provides coverage of 37,492 methylation probes with the large majority selected from conserved genomic loci across mammalian species and the remaining approximately two thousand selected based on known human biomarkers. Each probe contains a 50bp sequence on one side of a CpG site^12^. The methylation value of each probe is the beta value derived using SeSaMe normalization^36^ and represents the percent methylation.

### Definition of probe subset used for highest-coverage probe analyses

While most probes on the mammalian methylation array were selected from overall highly conserved genomic regions, for any given probe there could be non-human mammals for which it is not expected to work because of sequence divergence in that mammal or the sequence is non-unique in the mammal. To determine if a probe is expected to work in a mammalian species that has a genome available, we obtained previously computed mappability information^12^, which is whether the probe maps to a unique genomic location in that particular species. We obtained this information from annotation files at https://github.com/shorvath/MammalianMethylationConsortium/. In total, mappability information was available for 58 of the species included in the cross-validation analysis (Supplementary Data 4). We defined a subset of 11,749 probes as “highest-coverage” probes based on being mappable in at least 90% of these 58 species.

### CMImpute inputs and outputs for training and imputation

For training, CMImpute takes as input two matrices and outputs a single matrix. One of these input matrices is a *N_TRAIN_*x*M* matrix of methylation values where *N_TRAIN_* is the number of individual samples in the training dataset, *M* is the number of mammalian methylation array probes, the rows correspond to individuals, and the columns correspond to methylation probes. The other of these input matrices is a *N_TRAIN_*x*L* labels matrix that contains one-hot encoded species and tissue labels, where *L* is the sum of the total number of available species and tissues and the rows correspond to an individual while the columns correspond to either a species or a tissue. This *N_TRAIN_*x*L* matrix is a column-wise concatenation of an *N_TRAIN_*x*S* array of one-hot encoded species labels and an *N_TRAIN_*x*T* array of one-hot encoded tissue labels, where *S* is the number of species with observed data and *T* is the number of tissues with observed data. During training, CMImpute transforms the input into a *N_TRAIN_*x*Z* latent space representation where *Z* is the latent space dimension. The output is a *N_TRAIN_*x*M* matrix of reconstructed methylation samples, which contain CMImpute’s predicted methylation values of the original samples after reducing the original samples to a latent space representation. As with the input matrix of methylation values, the rows correspond to individuals and the columns correspond to methylation probes. During imputation, CMImpute takes as input just a 1x*L* labels matrix and outputs a 1x*M* matrix containing species-tissue combination mean sample values, where each value represents the average methylation value of a probe across individuals for a given species-tissue combination. The row corresponds to an imputed species-tissue combination and columns correspond to methylation probe values.

### CMImpute model

CMImpute uses a CVAE which is formally defined as a conditional directed graphical generative model and in practice is implemented as a neural network architecture that consists of an encoder, latent space, and decoder. The parameters of this architecture are trained to maximize the conditional log-likelihood^34^. The values the model components take on are dependent on the input observation matrix *X* and the corresponding label matrix *y*. The encoder representation is a recognition network *Q*_ϕ_ (*z*|*X*, *y*) which is used to approximate the true conditional prior *P*_θ_ (*z*|*y*), while the decoder representation is a generation network *P*_θ_ (*X*|*z*, *y*) where *z* denotes the encoded latent space vector.

In order to maximize the conditional log-likelihood *logP*(*X*|*y*), the theoretical variational lower bound is used as the objective function. The variational lower bound represents the lower bound for the probability of observations given the CVAE’s learned parameters and is given as 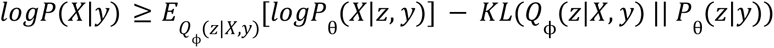^34^ where KL refers to the Kullback-Leibler divergence. The model assumes that both the encoder *Q*_ϕ_ (*z*|*X*, *y*) and true conditional prior *P*_θ_ (*z*|*y*) are multivariate Gaussians, where the learned latent space *z* is a vector sampled from *Q*ϕ (*z*|*X*, *y*) ∼ *N*(µ, Σ) where Σ is assumed to be diagonal and *P*_θ_ (*z*|*y*) ∼ *N*(0, *I*)^33,34,37^ where *I* is the identity matrix. Thus, the expectation term of the variational lower bound represents the expected output of the generation network and the KL term regularizes the latent space to be as close to *N*(0, *I*) as possible.

CMImpute represents the encoder and decoder as fully connected neural networks. The encoder takes as input training methylation samples *X* concatenated with their corresponding one-hot encoded species and tissue labels *y*. Each training sample in *X* is a vector of numbers between 0 and 1. With these inputs, the encoder predicts the latent space representation *z* based on the Gaussian parameters.

According to the variational lower bound and the definition of the encoder as the recognition network *Q*, *z* should be directly sampled from *Q*_ϕ_ (*z*|*X*, *y*). However sampling *z* from *Q* is a non-continuous operation, thus a gradient cannot be calculated and backpropagation cannot be used to learn the parameters if this sampling operation is performed within the network^37^. Instead the sampling operation is performed outside the network by using the “reparametrization trick” which reparametrizes the latent space, *z*, with a deterministic function *g*ϕ (*X*, *y*, ϵ) where ϵ ∼ *N*(0, *I*) is an auxiliary variable. This allows for the expectation term of the variational lower bound, 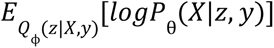, to be estimated via Monte Carlo (MC) approximation. By substituting this term with the approximation consisting of replacing *z* with the deterministic function *g*, the variational lower bound is replaced with a differentiable estimator in the form of the empirical lower bound:

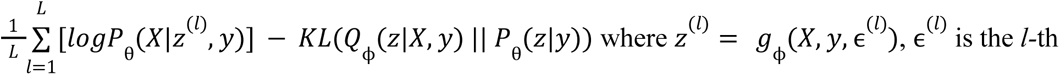

sample from the standard normal distribution, and *L* is the number of MC samples^33,34^. Following previous work, we set *L* to 1, which has been shown to yield an effective approximation in most settings^33^. As typical, μ and Σ were each represented as network layers directly before *z* in the encoder (Fig. 1b). μ was a vector representing the multivariate Gaussian distribution mean and σ was a vector representing the logarithms of each term of the Σ diagonal [σ = *log*(*diag*(Σ))].

Representing σ as a logarithm of the diagonal allows for an exponentiation operation during reparameterization, which makes computing the loss function (defined below) more numerically stable. Based on the “reparametrization trick” described above and the definitions of μ and σ, the latent space is defined as follows to approximate the true distribution while remaining differentiable: *z* = µ + *exp*(σ/2)ϵ where ϵ ∼ *N*(0, *I*)^37^ where *exp* denotes an operation that takes the exponentiation of each term of a vector.

The decoder takes as input *z* and *y* and attempts to reconstruct the training samples; the reconstructed samples are represented as *X’*. The training loss function (L_TRAIN_) represents the empirical lower bound via the sum of the reconstruction loss (binary cross entropy, L_RECON_) and KL regularization (L_REG_)^37^ :

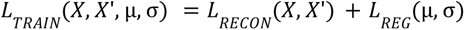

where

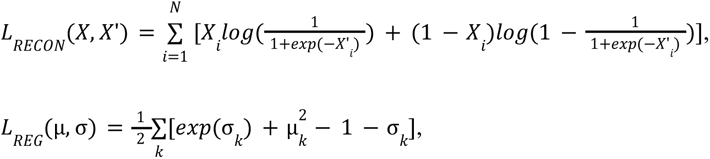

and *X_i_*is the *i*th training sample, *N* is the number of samples, and *k* is the latent space dimension.

### CMImpute training and hyperparameter selection

CMImpute was implemented in python 3.9.13 with Keras 2.10.0 (built on top of TensorFlow 2.10.0). CMImpute uses the Adam optimizer^38^ to learn the CVAE parameters: CMImpute selects the hyperparameters of the number of hidden layers in the encoder and decoder, hidden layer dimensions, activation function, latent space dimension, learning rate, and epsilon value via grid search (Supplementary Table 2, Supplementary Fig. 22). For each hyperparameter combination evaluated, a model was trained on a training dataset and used to impute combination mean samples for species and tissue combinations in a corresponding validation dataset. Validation datasets consisted of species and tissue combinations not present in the training dataset, but where at least one same-species and same-tissue sample was present in the training dataset. CMImpute then selected the hyperparameter combination with the highest average sample-wise Pearson correlation between the imputed and held-out species-tissue combinations in the validation dataset.

### Species-tissue combination mean imputation

Species-tissue combination mean imputation refers to using a trained model to predict a sample that represents a species’ average methylation values in a specific tissue. CMImpute uses its trained decoders to generate species-tissue combination mean samples for every desired combination of species and tissues via the following steps (Fig. 1b):

1. CMImpute draws a random sample from a standard normal distribution of shape 1x*Z* where *Z* is the latent space dimension. This sample is used as the latent space representation. CMImpute performed this random normal sampling with Numpy version 1.23.4.
2. CMImpute inputs the random normal latent space representation from step 1 and a 1x(*S*+*T*) one-hot encoded label of the target species and tissue type into the decoder, where *S* and *T* are the number of profiled species and tissue types, respectively. The resulting output of the decoder is an imputed species-tissue combination mean sample for the target combination.

In previous CVAE applications, the generative process is composed of two parts: i) obtaining a latent representation by inputting a chosen input into the encoder and ii) generating a new sample by inputting the resulting latent representation along with a conditional label into the decoder^30,32,34,39,40^. Using an encoded input as the latent representation makes the resulting generated output dependent on the chosen input sample and thus specific to the individual from which the sample was obtained. However, for the problem of imputing species-tissue combination mean samples that we consider here, an individual-agnostic generative process based on the species and tissue label is required. Since the provided conditional label is sufficient to drive sample imputation of a specific species and tissue combination^32,34^ and the latent space is a multivariate Gaussian regularized to be close to a standard normal distribution, CMImpute samples from a standard normal distribution to obtain a latent representation. CMImpute’s sampling scheme approximates a true sampling from the latent space, as the latent space is regularized to be as close to the standard normal distribution as possible, without encoding information from existing methylation samples. We verified the specific sampled latent space values have minimal impact on the final predicted sample (Supplementary Fig. 23), and thus the imputed samples are mainly based on the overall species and tissue label inputted into the trained decoder.

### Logistic regression baseline

We compared CMImpute to a logistic regression baseline with L_2_ regularization. For this baseline, we trained one model per methylation probe with separate species and tissue features. For a particular probe and species and tissue combination, the trained model was then used to predict the methylation value. Specifically the predicted value for a probe was: 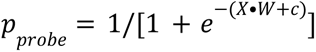, where *X* was the one-hot-encoded representation of the species and tissue labels, *W* the learned feature weights, and *c* the learned intercept. The feature weights were learned using the loss function *L*_*LOGLOSS*_ =− *ylog*(*p*_*probe*_) − (1 − *y*)*log*(*p*_*probe*_) + λ‖*W*‖^2^ where *y* is the real methylation probe values and λ was the regularization coefficient. We trained each logistic regression model in python3.9.13 using scikit-learn version 0.24.2. Using this package, we included each one-hot-encoded species and tissue label from the training dataset twice in the training input, once with a corresponding y-label of 1 and sample weight of the probe’s methylation value and once with a corresponding y-label of 0 and sample weight of one minus the probe’s methylation value. In this setup, the methylation value prediction corresponds to the probability of a positive classification. Once trained, we concatenated the predictions of each probe-specific model together to form a full imputed methylation sample for a particular species and tissue combination.

For the cross-validation analysis, we tuned the regularization coefficient across λ values of 1, 2, 4, 8, and 16. We selected λ that yielded combination mean samples with the highest Pearson correlation with held-out samples. A λ value of 2 yielded this highest testing performance (0.886 Pearson correlation, Supplementary Fig. 24).

### Mean imputation baselines

We compared CMImpute to three mean imputation baselines referred to as the species baseline, tissue baseline, and global baseline. Let *N* be the total number of experimentally profiled samples, *X_i_* an individual methylation sample (1x*M* where *M* is the total number of probes), and *X*’_*S*,*T*_ the imputed combination mean sample representing the species *S* and tissue *T*.

1) The species baseline imputed a combination mean sample by taking the average of all training samples of the target species. 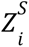 is an indicator variable indicating whether a sample *i* is from a particular species *S*, and *N_S_* is the number of experimentally profiled samples within a species *S*.

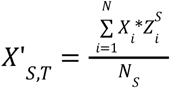

2) The tissue baseline imputed a combination mean sample by taking the average of all training samples of a target tissue. 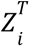 is an indicator variable indicating whether a sample *i* is from a particular tissue *T*, and *N_T_* is the number of experimentally profiled samples within a tissue *T*.

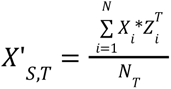

3) The global baseline imputed a combination mean sample by taking the average of all training samples.

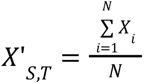

### Cross-validation datasets to compare imputed species-tissue combination mean samples with held-out observed data

To compare CMImpute and baseline predictions to held-out observed species-tissue combination mean samples, we created multiple training and testing datasets. We considered the 520 observed species-tissue combinations where the target species is available in more than one tissue type and the target tissue is available in more than one species. We randomly divided these combinations into five folds, resulting in 465 imputed species-tissue combinations for evaluations. These 465 combinations correspond to 134 species with data from more than one tissue type available and 23 tissues with data from more than one species available. This final amount is less than the 520 combinations initially considered because we only considered the imputation performance of a species-tissue combination if there was both same-species different-tissue and same-tissue different-species data available in the corresponding training dataset.

In cross-validation, we considered each of the five folds a testing dataset. When each fold was considered as a testing dataset, the remaining data outside the fold was included in either the training or validation datasets. To determine which combinations were included in the training or validation dataset, we first randomly divided the combinations into candidate training and validation datasets. To perform this division, we randomly selected 20% of the species-tissue combinations to form the candidate validation dataset, while the remaining 80% of combinations formed the candidate training dataset. For each combination in the candidate validation dataset, if the combination did not have at least one combination of the same species and at least one combination of the same tissue present in the candidate training dataset, then the combination was moved from the candidate validation dataset to the candidate training dataset. If at the end of this procedure the candidate validation dataset consisted of less than 10% of the remaining combinations outside the testing fold, we made a new split of the training and validation data, and repeated the process. Otherwise, the candidate training and validation datasets were used for the training and validation datasets. This process resulted in the validation dataset consisting of at least 10% of the combinations remaining outside the testing fold while still having same-species and same-tissue information available in the training dataset.

The hyperparameters (activation function, latent space dimension, learning rate, epsilon value, number of hidden layers, and hidden layer dimensions) were selected via grid search based on the validation performance. Using the models selected based on hyper-parameter tuning we imputed species-tissue combination mean samples representing combinations held-out from the corresponding training and validation sets.

For performance evaluations, we concatenated all imputed samples into one grid of species-tissue combination mean samples. For methylation value and pairwise correlation visualizations, we ordered both the samples and probes based on hierarchical clustering followed by optimal leaf ordering^35^.

### Prediction of non-observed species-tissue combinations

To impute non-observed species and tissue combinations, we first selected the hyperparameters for a model. We did this by creating four random 80%-20% training-testing splits on observed combinations that involve a species with more than one tissue type and a tissue type with more than one species available. This criteria ensures that when a combination is held-out during hyperparameter tuning, same-species different-tissue and same-tissue different-species training information will still be available during training. Of the 746 experimentally profiled species-tissue combinations, 520 combinations satisfied this criteria (Supplementary Data 1). Once created, we performed a hyperparameter grid search on each of the four splits and determined the highest performing hyperparameter combination for each split based on sample-wise Pearson correlation with held-out samples. For each of these four best hyperparameter combinations, we averaged the performance for those hyperparameters across all four random splits and selected the hyperparameters that resulted in the highest average. We saw the highest sample-wise performance on average across all tuning datasets (0.933) for the following hyperparameter combination: two hidden layers of dimensions 1024 and 512, TanH activation function, latent space dimension of 8, learning rate of 0.001, and epsilon value of 0.0001. We then trained a single model based on these hyperparameter values using all available methylation samples. Finally we used the trained model to generate samples of species-tissue combinations not experimentally profiled.

### Probe variance calculations

We measured three types of variances across the experimentally profiled data to determine how a probe’s variance among different species and tissues impacts imputation performance. These three types of probe-wise variances are: (1) inter-combination variance which measures the variance between species and tissue combinations, (2) mean inter-tissue variance which measures the average variance between tissues within a species, and (3) mean inter-species variance which measures the average variance between species within a tissue type.

Let *M* be the number of probes in the mammalian methylation array. We used the following process for calculating the probe-wise inter-combination variance 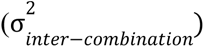 (1x*M* vector) of the experimentally profiled data:

1) Calculate the mean methylation of each observed species and tissue combination (e.g. human heart, horse liver, etc.). This step prevents the number of individual samples in a particular combination from skewing the variance calculation so the variance is measured between combination mean samples. In the equations below, let *N* be the number of observed samples, *N*_*species,tissue*_ be the number of observed samples within a species and tissue combination, *X_i_* be an individual methylation sample (1x*M*), 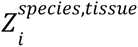 indicator variable indicating whether a sample *i* is from a particular species and tissue combination, *X*’_*species,tissue*_ be the resulting mean methylation of a particular species and tissue combination (1x*M*), *C* be the set of observed species and tissue combinations, and *X*’_*combos*_ be the resulting |*C*|x*M* array containing the mean methylation value of each probe for each unique species-tissue combination. Formally *X*’_*species,tissue*_ and *X*’_*combos*_ are defined as

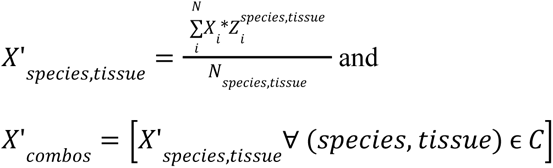

2) Calculate the variance of each probe across each unique species-tissue combination.

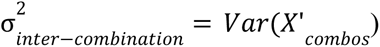

Below is the process for calculating the probe-wise mean inter-tissue variance 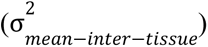 (1x*M* vector) of the observed data.

1) Let *S* represent the set of species with more than one tissue type available within the species. Let *X* represent the individual methylation samples from all species in *S*.
2) For each species in *S*, calculate the mean methylation of each tissue. Let *T*_*species*_ be the set of observed tissues within a species. *X*’_*species*_ is the resulting |*T*_*species*_|x*M* array containing the average methylation value for each tissue observed in the target species. Formally *X*’_*species*_ is defined as

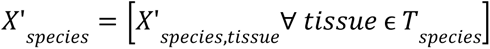

3) For each species in *S*, calculate the variance of each probe across each tissue available in the species. 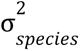 is the resulting variance of each probe across each tissue observed in the target species, that is

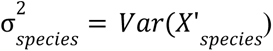

4) Calculate the average variance for each probe across all species in *S*.

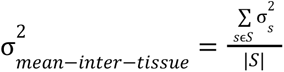

Below is the process for calculating the probe-wise mean inter-species variance 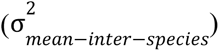 (1x*M* vector) of the observed data.

1) Let *T* represent the set of tissues profiled in more than one species. Let *X* represent the individual methylation samples from all tissues in *T*.
2) For each tissue in *T*, calculate the mean methylation of each species. *S*_*tissue*_ is the set of experimentally profiled species within a tissue. *X*’_*tissue*_is the resulting |*S*_*tissue*_|x*M* array containing the average methylation value for each tissue observed in the target species. Formally *X*’_*tissue*_ is defined as

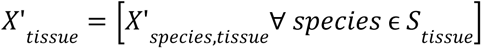

1) For each tissue in *T*, calculate the variance of each probe across each species available in the tissue. 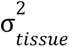 is the resulting variance of each probe across each species observed in the target tissue, that is

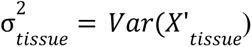

2) Calculate the average variance for each probe across all tissues in *T*.

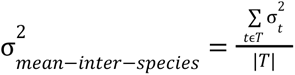

### Linear regression analysis of species-tissue combination mean samples relative to species maximum lifespan

We evaluated how predictive combination mean methylation samples were of the log-maximum lifespan of a species in a linear regression model through a leave-one-species-out (LOSO) analysis. For this we used maximum lifespan values for 114 species obtained from the anAge database^41^. Our methodology follows a similar structure of Li et al^16^. We implemented and trained the linear regression models in python 3.9.13 using scikit-learn version 0.24.2. The LOSO analysis was performed in the following four settings:

1. Tissue-agnostic observed samples: Observed species-tissue combination mean samples for each observed tissue and species combination were averaged across all tissue types within a species to form a single average observation per species. The number of tissues being averaged in each species sample is dependent on the number of observed tissues available.
2. Tissue-agnostic imputed samples: Imputed species-tissue combination mean samples for each non-observed species and tissue combination were averaged across all tissue types within a species to form a single average sample per species. The number of tissues being averaged in each species sample is dependent on the number of imputed tissues available.
3. Tissue-specific observed samples: Instead of averaging across tissue types, each observed species-tissue combination mean sample remains intact and was used as a training sample. Each tissue within a species shares the same log-maximum lifespan. The number of samples held out in the LOSO analysis corresponds to the number of tissue types observed in the species.
4. Tissue-specific imputed samples: Imputed species-tissue combination mean samples were used as training samples. Each tissue within a species shares the same log-maximum lifespan. The number of samples held out in the LOSO analysis corresponds to the number of tissue types that are not observed in the species. The imputed combination mean samples span the same 114 species as the observed setting, but the tissues represented in a species do not overlap with the observed samples.

We evaluated the predictive performance of the tissue-agnostic species’ averages for both the observed and imputed data by computing the Pearson correlation and MSE between the predicted and reported log-maximum lifespans. We also calculated the Pearson correlation between the imputed and observed predicted log-maximum lifespan values across all species. We similarly evaluated the predictive performance of the tissue-specific species-tissue combination mean samples for both observed and imputed data using Pearson correlation and MSE. For each tissue, we computed the Pearson correlation and MSE between the predicted log-maximum lifespan for each species in which the tissue was observed and the reported log-maximum lifespan and averaged the Pearson correlations and MSEs across the tissues. We restricted this analysis to tissue types observed in three or more species as the Pearson correlation between samples of length one or two is not valid.

## Data and code availability

CMImpute code and the full grid of imputed species-tissue combination mean samples can be found at https://github.com/ernstlab/CMImpute. All data used was previously published by the Mammalian Methylation Consortium^10^ and available from the Gene Expression Omnibus GSE223748. Genome annotations can be found at https://github.com/shorvath/MammalianMethylationConsortium.

## Acknowledgements

We thank the Mammalian Methylation Consortium for generating the mammalian methylation array data. We thank Caesar Li for assistance with using the data. This work was supported by the the National Institutes of Health (NIH) (DP1DA044371, U01MH130995) (J.E.), UCLA Jonsson Comprehensive Cancer Center, Eli and Edythe Broad Center of Regenerative Medicine and Stem Cell Research Ablon Scholars Program (J.E.), and the NIH Training Grant in Genomic Analysis and Interpretation T32HG002536 (E.M.).

## Author Contributions

E.M. and J.E. developed the CMImpute method. E.M. implemented the method, applied it to the mammalian methylation array data and performed analyses. J.E. conceived the study and supervised the project. S.H. provided data and advice for the project. E.M. and J.E. wrote the main text. All authors participated in editing the text.

