## Supplementary Figures for "Cross-species and tissue imputation of species-level DNA methylation samples across mammalian species"

**Contents**

Supplementary Figures 1-24
Supplementary Tables 1-2

### Supplementary Figures

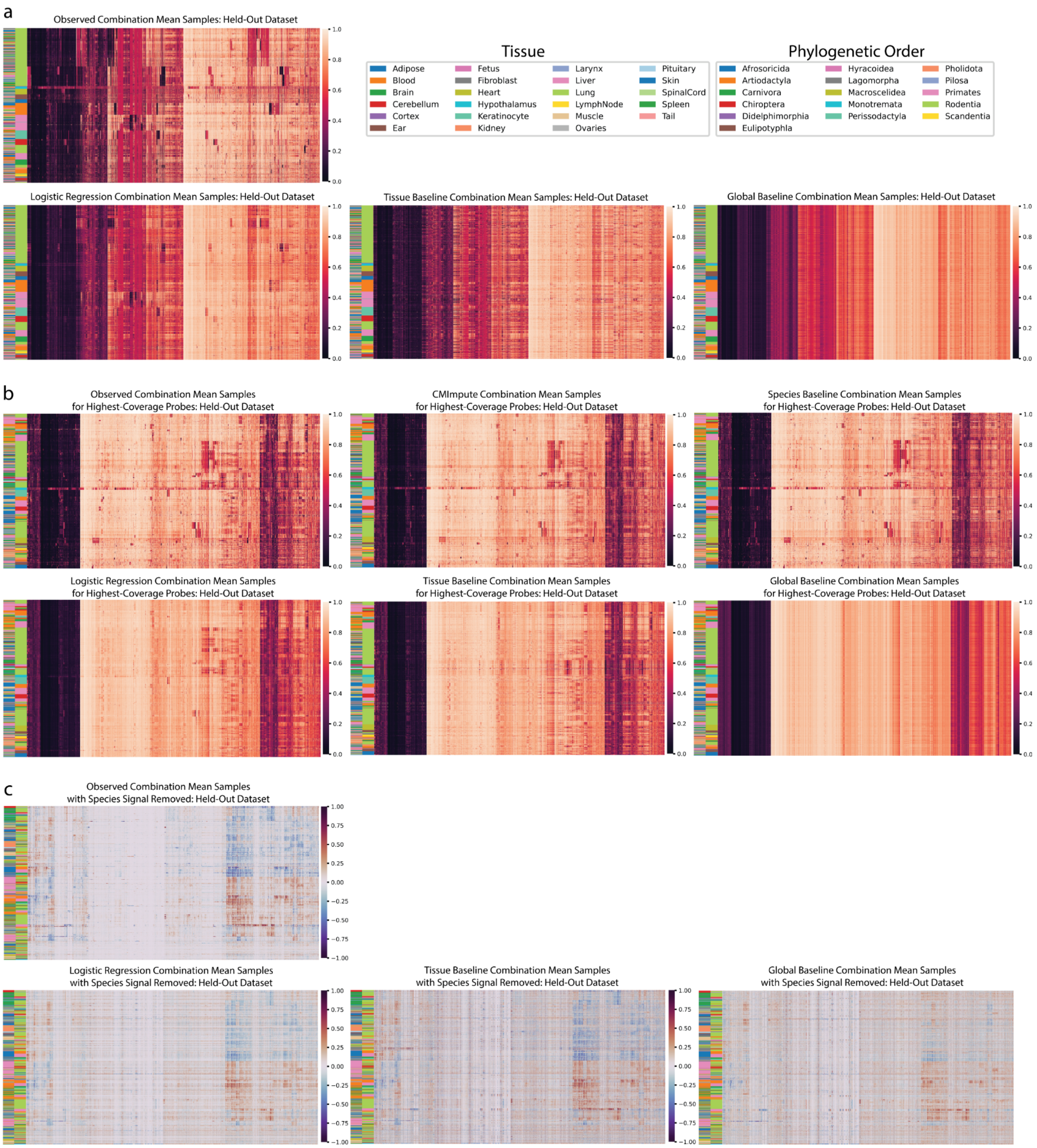


**Supplementary Figure 1. Imputed species-tissue combination mean sample visualizations.**

**a)** Similar heatmaps as in Fig. 2a-c of predictions of held-out datasets' methylation probe values shown here for logistic regression, tissue baseline, and overall baseline (bottom row; left to right). Observed methylation probe values included again for comparison (top row). Each row is a species-tissue combination mean sample and each column is a methylation probe. Samples and probes were ordered based on hierarchical clustering followed by optimal leaf ordering. Color bars on the left indicate the phylogenetic order (inner) and tissue (outer) corresponding to the samples. Legends corresponding to the color bars can be found in the top right of the figure. Color scale representing methylation values from 0 to 1 on the right. **b)** Heatmaps of methylation probe values for the observed data held-out during cross-validation and CMImpute and baseline predictions restricted to the highest coverage probes. Samples and probes are ordered and labeled similarly to a. **c)** Similar heatmaps as in Fig. 2d-e of predicted datasets with the species signal removed shown here to highlight the differentially methylated tissue regions for logistic regression, tissue baseline, and overall baseline. Observed dataset with species signal removed included again for comparison. Color scale representing methylation delta values from -1 to 1 on the right.

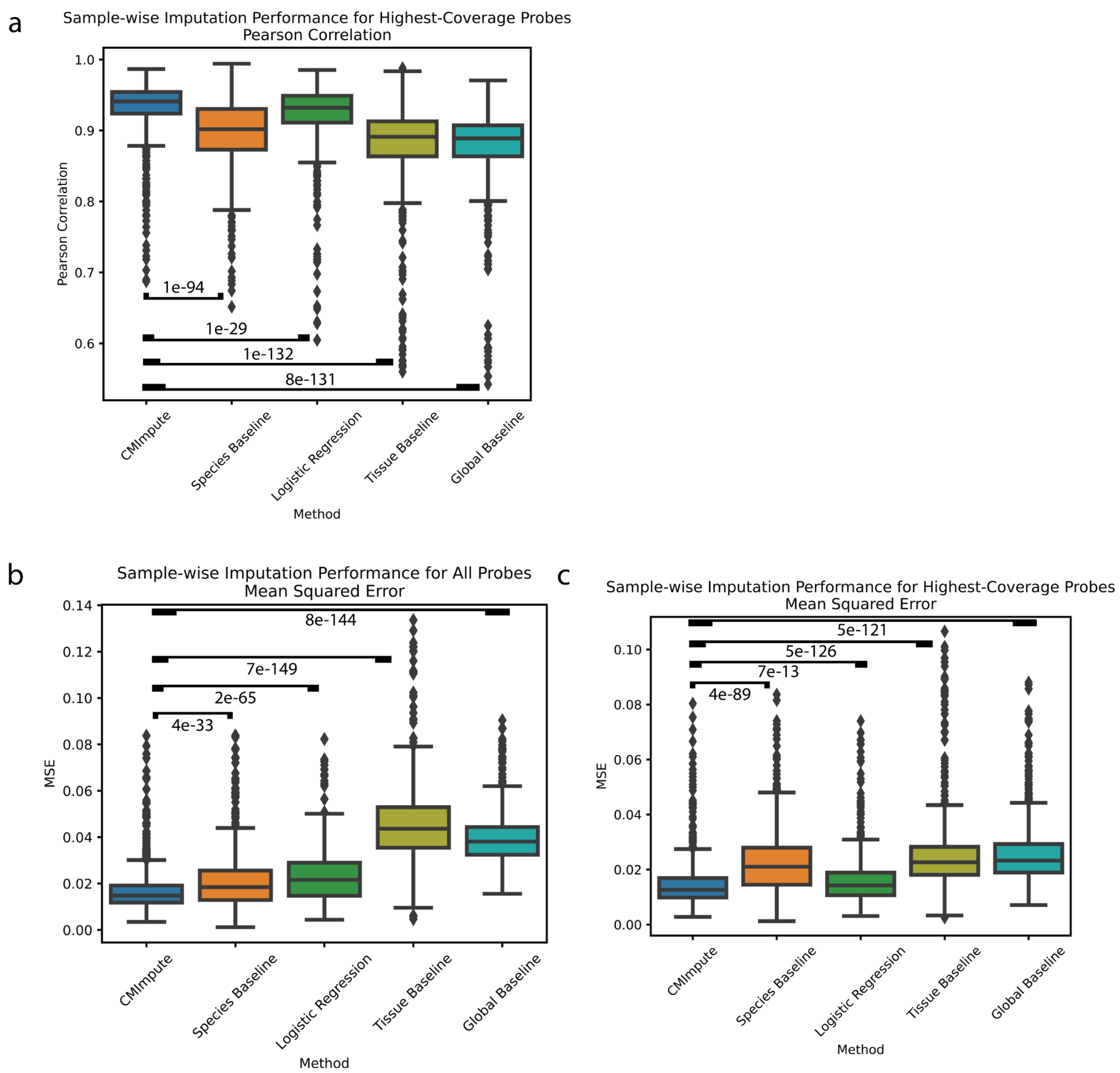


**Supplementary Figure 2. Sample-wise performance distributions. a)** Box-plot showing distribution of sample-wise Pearson correlation of imputed species-tissue combination mean samples with held-out observed values when restricted to highest-coverage methylation probes for CMImpute and all baselines. Baselines labeled by Wilcoxon signed-rank test p-value comparing CMImpute's sample-wise Pearson correlation and each baseline's sample-wise Pearson correlation for each imputed combination ([CMImpute, Species Baseline], [CMImpute, Logistic Regression], [CMImpute, Tissue Baseline], [CMImpute, Overall Baseline]). **b-c)**

Sample-wise MSE of imputed combination mean samples based on **b)** all probes and **c)** highest-coverage probes only.

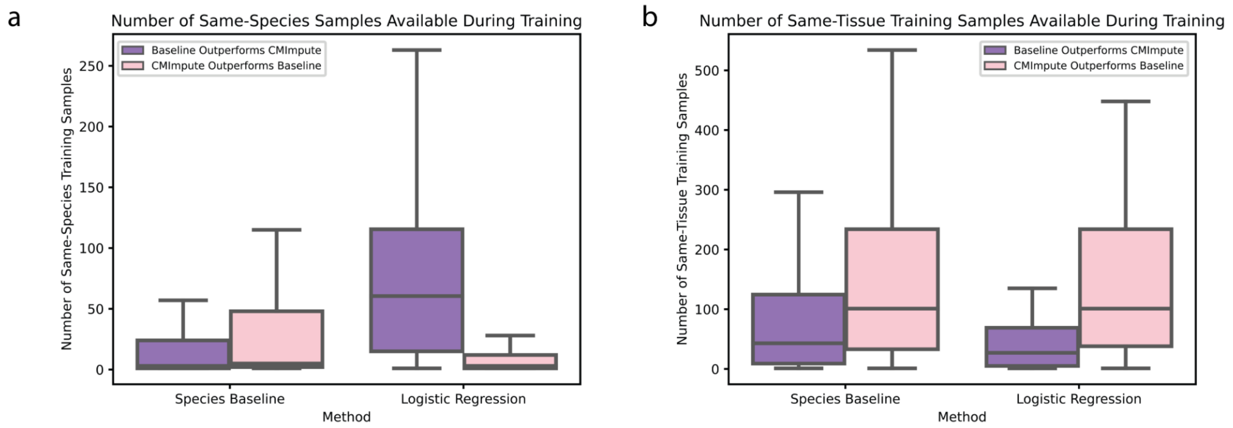


**Supplementary Figure 3. Analysis of subset of samples where species baseline and logistic regression outperform CMImpute. a-b)** The number of **a)** individual same-species samples and **b)** individual same-tissue samples available during model training for imputed combination mean samples where CMImpute outperforms the species baseline and logistic regression and vice versa. The imputed combination mean samples where the baseline outperforms CMImpute are the 32% of samples (291 out of 907 combination mean samples) below the red line in the top left plot of Fig. 3b for the species baseline and the 22% (200 out of 907 combination mean samples) of samples below the red line in the top right plot of Fig. 3b for logistic regression.

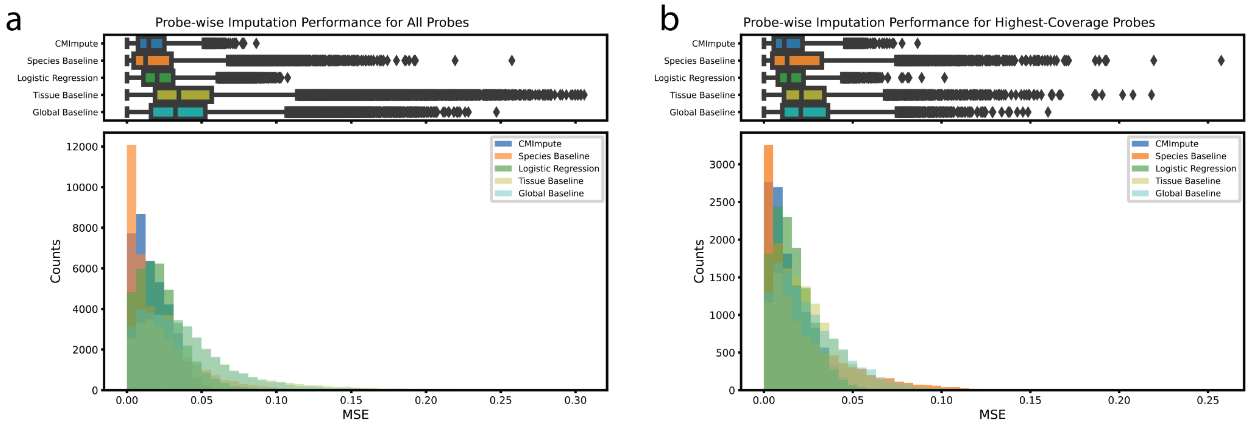


**Supplementary Figure 4. Probe-wise mean squared error distributions. a)** Distributions of probe-wise MSE with held-out observed values based on all probes for CMImpute and all baselines. Plots are formatted in the way as Fig. 4a-b. The top boxplots show the distribution of probe-wise correlations with held-out observed values. The bottom boxplots show the number of imputed combination mean samples across 50 performance bins. Legend for both the boxplot and histogram shown in histogram plot. CMImpute yields the best mean probe-wise MSE of 0.017 compared to the species baseline's 0.021, logistic regression's 0.023, the tissue baseline's 0.045, and the global baseline's 0.039. Corresponding plots for subsets of higher variance probes can be found in Supplementary Fig. 12d-f. **b)** Same as a except restricted to highest-coverage probes. CMImpute yields the best mean probe-wise MSE of 0.014 compared to the species baseline's 0.023, logistic regression's 0.016, the tissue baseline's 0.025, and the global baseline's 0.026. Corresponding plots for subsets of higher variance probes can be found in Supplementary Fig. 12a-c.

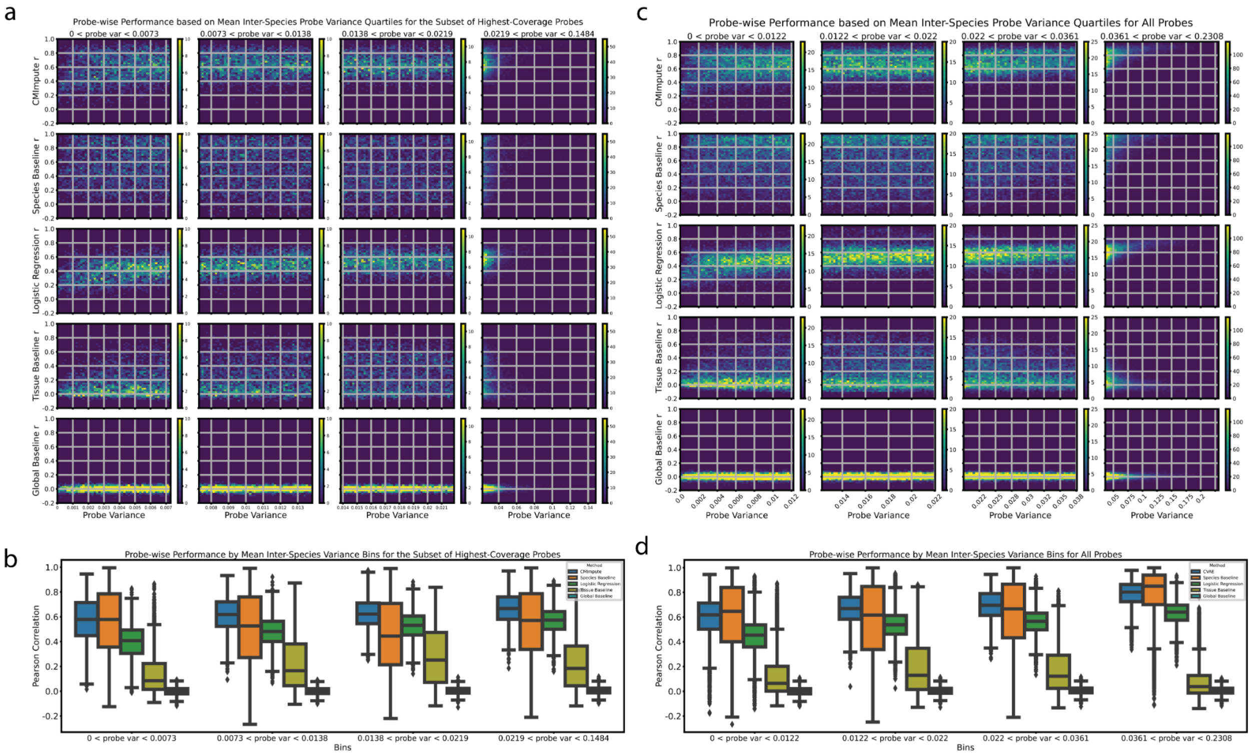


**Supplementary Figure 5. Probe-wise Pearson correlation relative to mean inter-species probe variance. a)** 2-d histograms comparing mean inter-species variance with probe-wise performance when considering the subset of highest-coverage probes with CMImpute, species baseline, logistic regression, tissue baseline, and global baseline (displayed from top to bottom) probe-wise Pearson correlation. Heatmaps are formatted in the same way as Fig. 4c-d. Each row contains four heatmaps corresponding to variance quartiles. Within each variance quartile, the heatmap shows the number of probes within a probe variance bin along the x-axis and a probe-wise Pearson correlation bin along the y-axis split into 50 bins along each axis. **b)** Boxplots of probe-wise Pearson correlation with held-out observed values for highest-coverage probes in each mean inter-species variance quartile. Each variance quartile represented in the boxplots correspond to the variance quartile in the 2-d histograms from a. **c)** Same as a) but when considering all probes. **d)** Same as b) but when considering all probes.

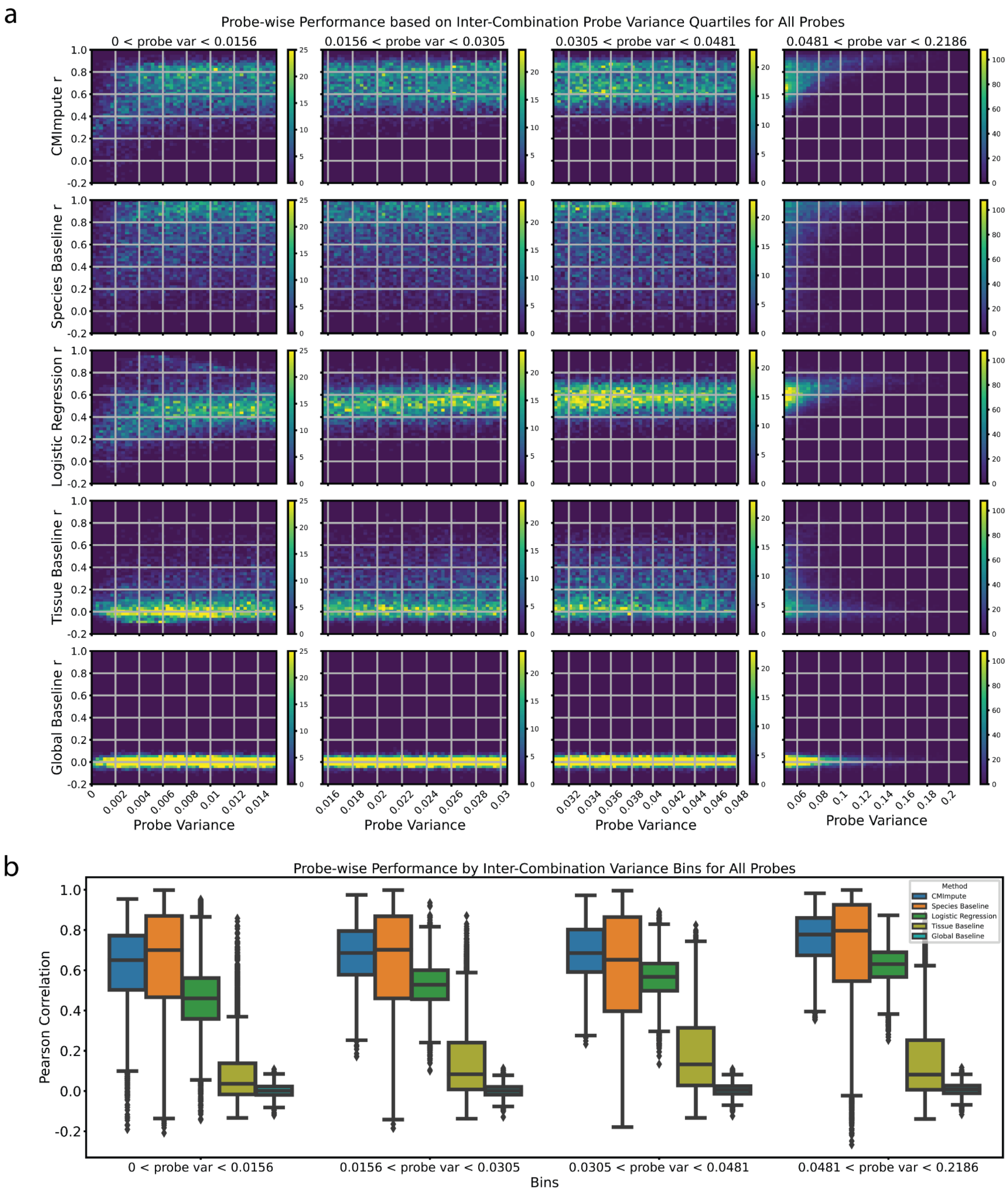


**Supplementary Figure 6. Probe-wise Pearson correlation relative to inter-combination probe variance. a)** 2-d histogram in the same format as Supplementary Fig. 5a,c but here comparing inter-combination variance with probe-wise performance when considering all probes. **b)** Boxplots of probe-wise Pearson correlation in the same format as Supplementary Fig. 5b,d for each inter-combination variance quartile when considering all probes.

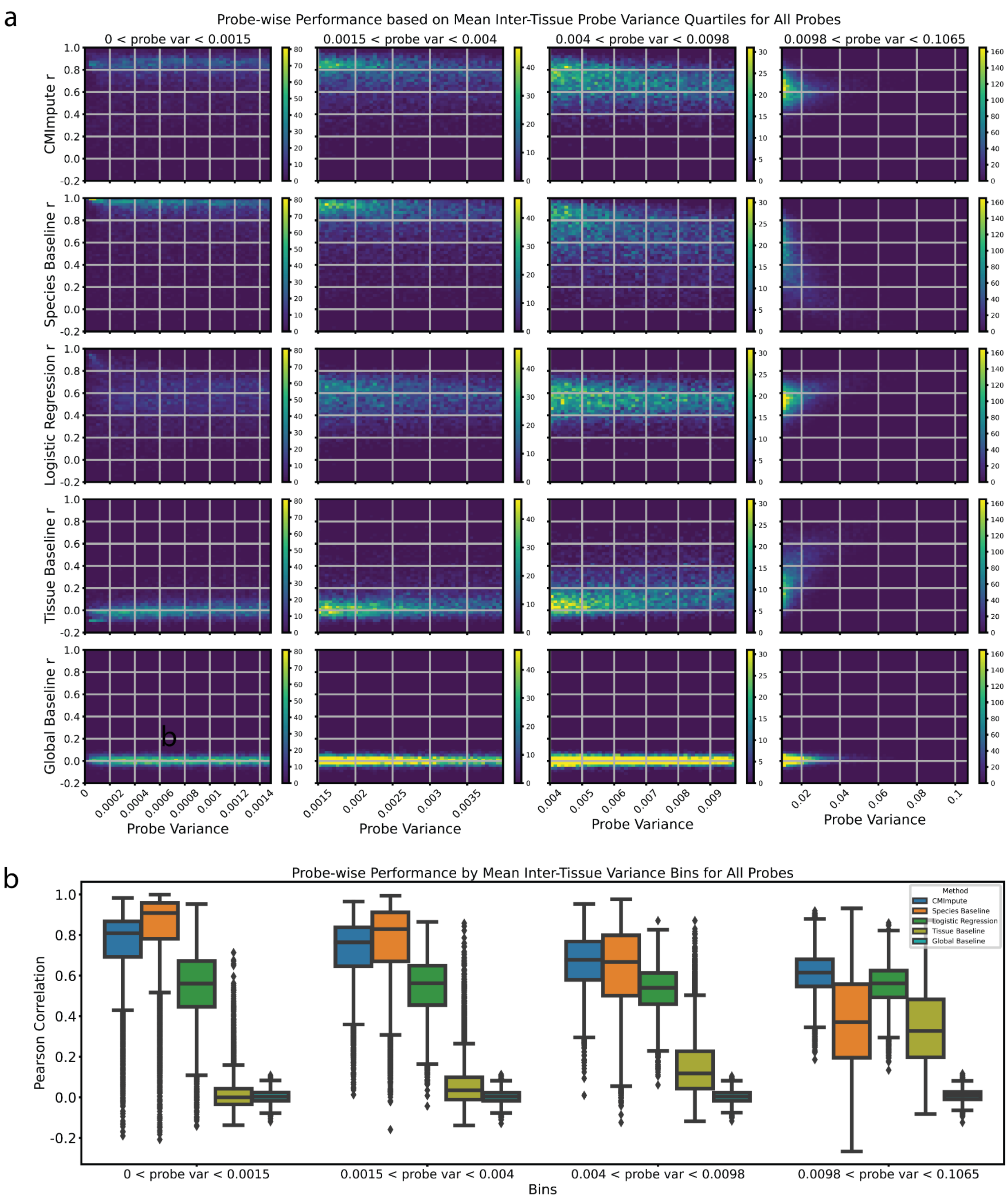


**Supplementary Figure 7. Probe-wise Pearson correlation relative to mean inter-tissue probe variance. a)** 2-d histograms comparing mean inter-tissue variance with CMImpute, species baseline, logistic regression, tissue baseline, and global baseline (displayed from top to bottom) probe-wise Pearson correlation when considering all probes. Heatmaps are formatted in the same way as Fig. 4c-d. Each row contains four heatmaps corresponding to variance quartiles. Within each variance quartile, the heatmap shows the number of probes within a probe variance bin along the x-axis and a probe-wise Pearson correlation bin along the y-axis split into 50 bins along each axis. **b)** Boxplots of probe-wise Pearson correlation with held-out observed values for probes in each mean inter-tissue variance quartile when considering all probes. Each variance quartile represented in the boxplots corresponds to the variance quartile in the 2-d histograms from a.

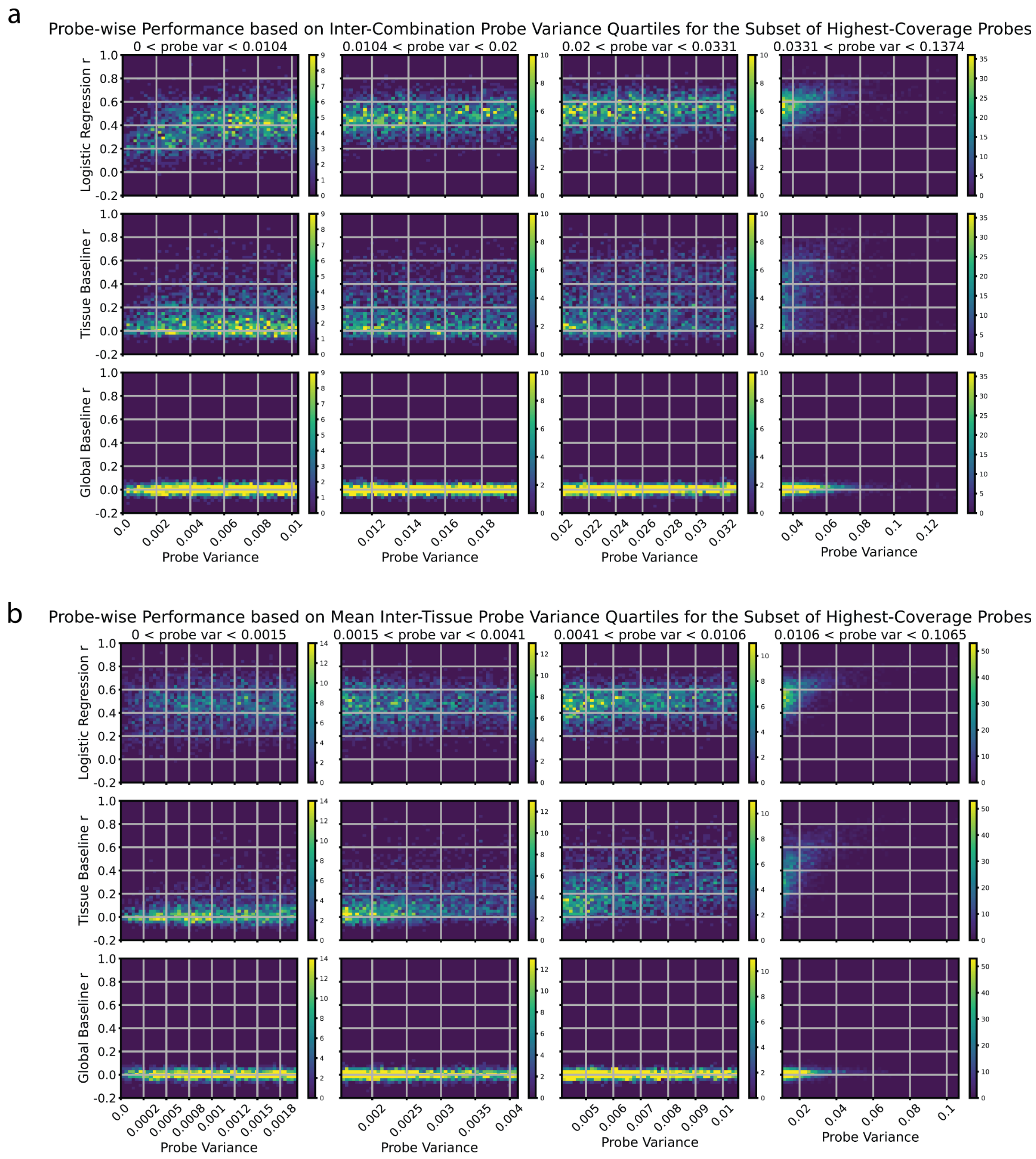


**Supplementary Figure 8. Probe-wise Pearson correlation relative to inter-combination and mean inter-tissue probe variance for baselines. a)** Similar 2-d histogram as Fig. 4d comparing inter-combination variance with probe-wise Pearson correlation shown here for logistic regression (top row), tissue baseline (middle row), and overall baseline (bottom row) probe performance when considering the subset of highest-coverage probes. **b)** Similar 2-d histogram as Fig. 4c comparing mean inter-tissue variance with probe-wise Pearson correlation shown here for logistic regression (top row), tissue baseline (middle row), and overall baseline (bottom row) probe performance.

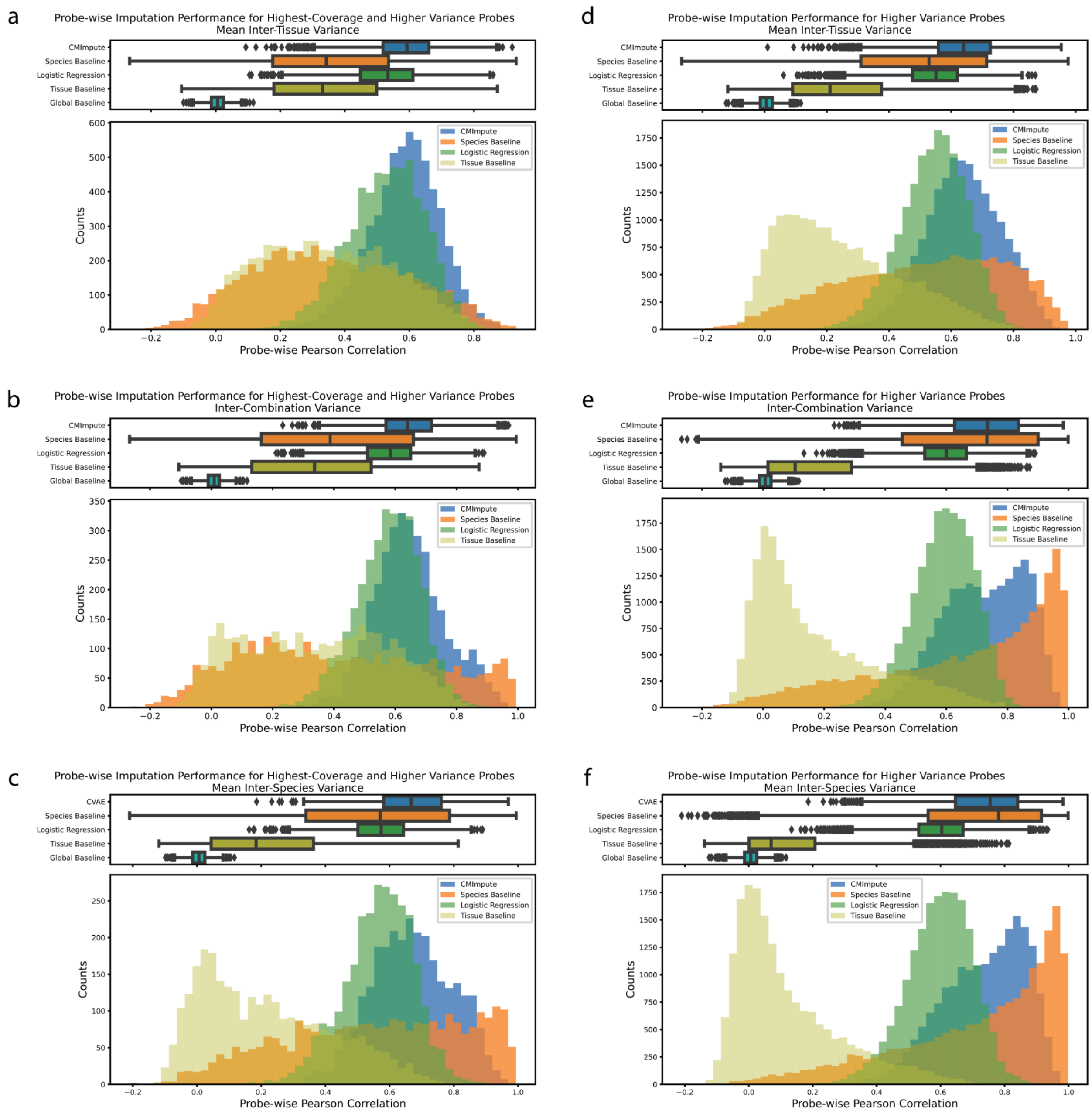


**Supplementary Figure 9. Probe-wise performance distributions for higher variance and highest-coverage probes.** Distributions of probe-wise Pearson correlations with held-out observed values for **a)** 5,997 probes that are among the set of the highest-coverage probes and have a mean inter-tissue variance > 0.004 (median mean inter-tissue variance among set of highest-coverage probes), **b)** 3,390 probes that are among the set of the highest-coverage probes and have an inter-combination variance > 0.031 (median inter-combination variance among set of highest-coverage probes), **c)** 2,883 probes that are among the set of the highest-coverage and have a mean inter-species variance > 0.022 (median mean inter-species variance among set of highest-coverage probes), **d)** 18,746 probes with a mean inter-tissue variance > 0.004 (median mean inter-tissue variance), **e)** 18,746 probes with an inter-combination variance > 0.031 (median inter-combination variance), and **f)** 18,746 probes with a mean inter-species variance > 0.022 (median mean inter-species variance). The top boxplots show the distribution of probe-wise correlations with held-out observed values. The bottom boxplots show the number of imputed combination mean samples across 50 performance bins. Legend for both boxplots and histograms shown in histogram plot and indicate which method (CMImpute and all baselines) are being considered.

a

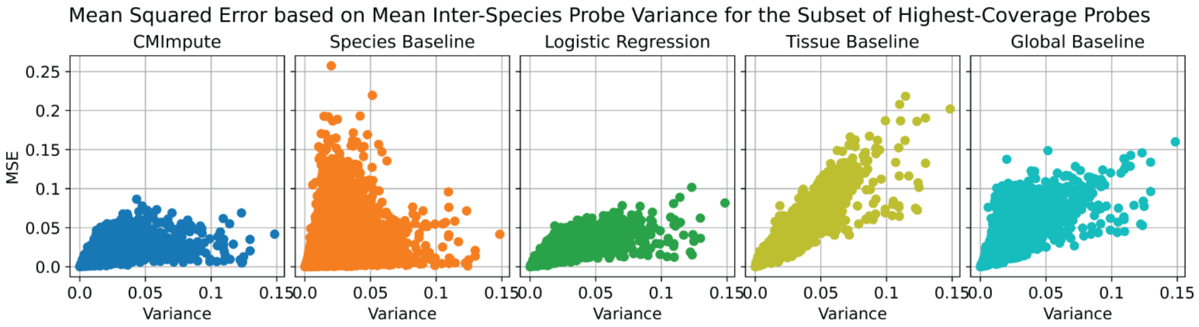
Mean Squared Error based on Mean Inter-Species Probe Variance for the Subset of Highest-Coverage Probes
CMImpute
Species Baseline
Logistic Regression
Tissue Baseline
Global Baseline
MSE
Variance

b

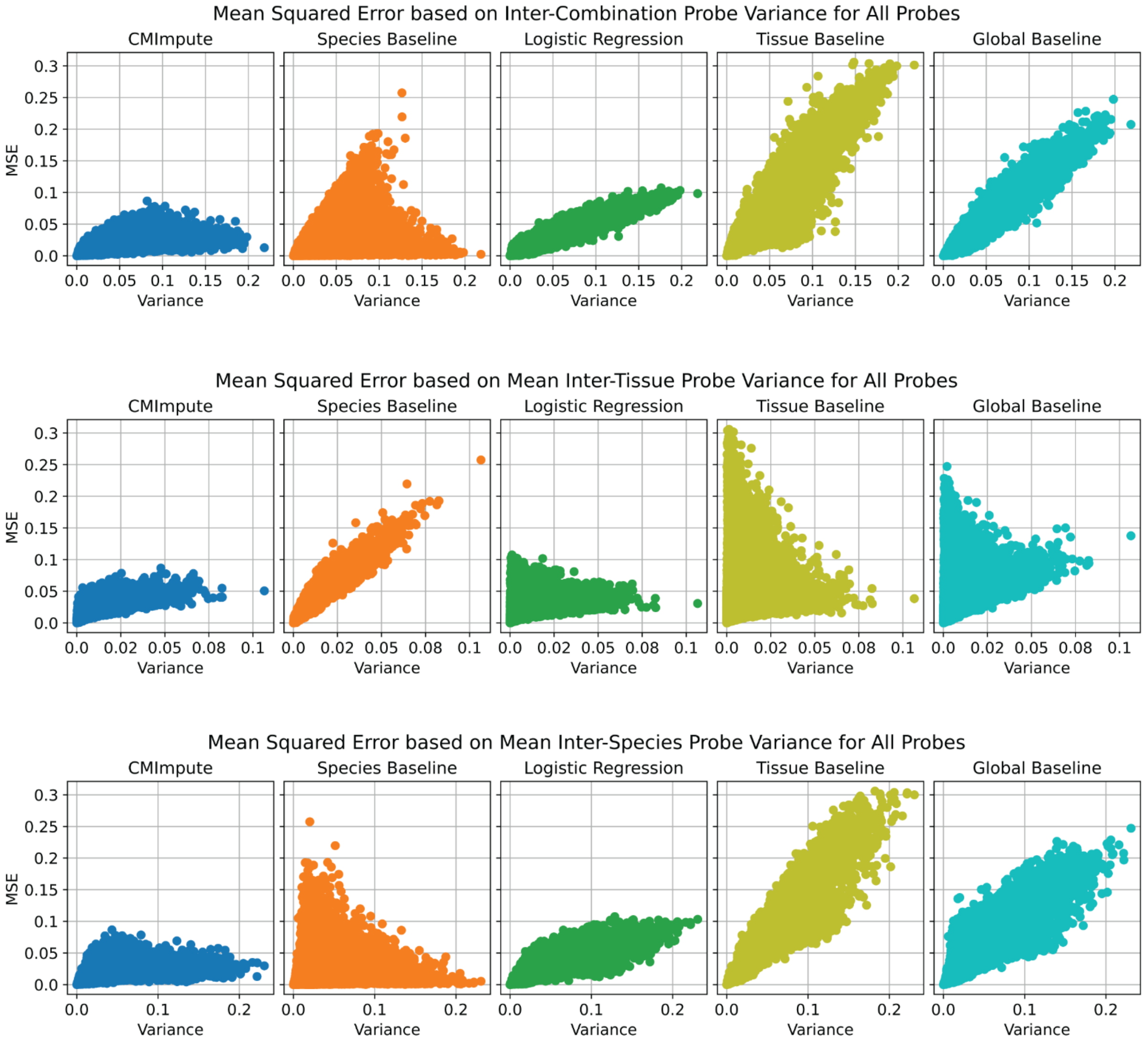
Mean Squared Error based on Inter-Combination Probe Variance for All Probes
Mean Squared Error based on Mean Inter-Tissue Probe Variance for All Probes
Mean Squared Error based on Mean Inter-Species Probe Variance for All Probes
CMImpute
Species Baseline
Logistic Regression
Tissue Baseline
Global Baseline
MSE
Variance

**Supplementary Figure 10. Probe-wise MSE relative to probe variance. a)** Scatterplots showing the relationship between probe-wise MSE and mean inter-species variance restricting to the highest-coverage probes in the same format as Fig. 4g-h. The y-axis is the probe-wise MSE with held-out observed values for CMImpute, species baseline, logistic regression, tissue baseline, or overall baseline (left to right). The x-axis is the probe variance. Each dot corresponds to a single imputed probe. **b)** Similar scatter plots to a) and Fig. 4g-h comparing the probe-wise MSE with inter-combination (top), mean inter-tissue (middle), and mean inter-species (bottom) variance when considering all probes.

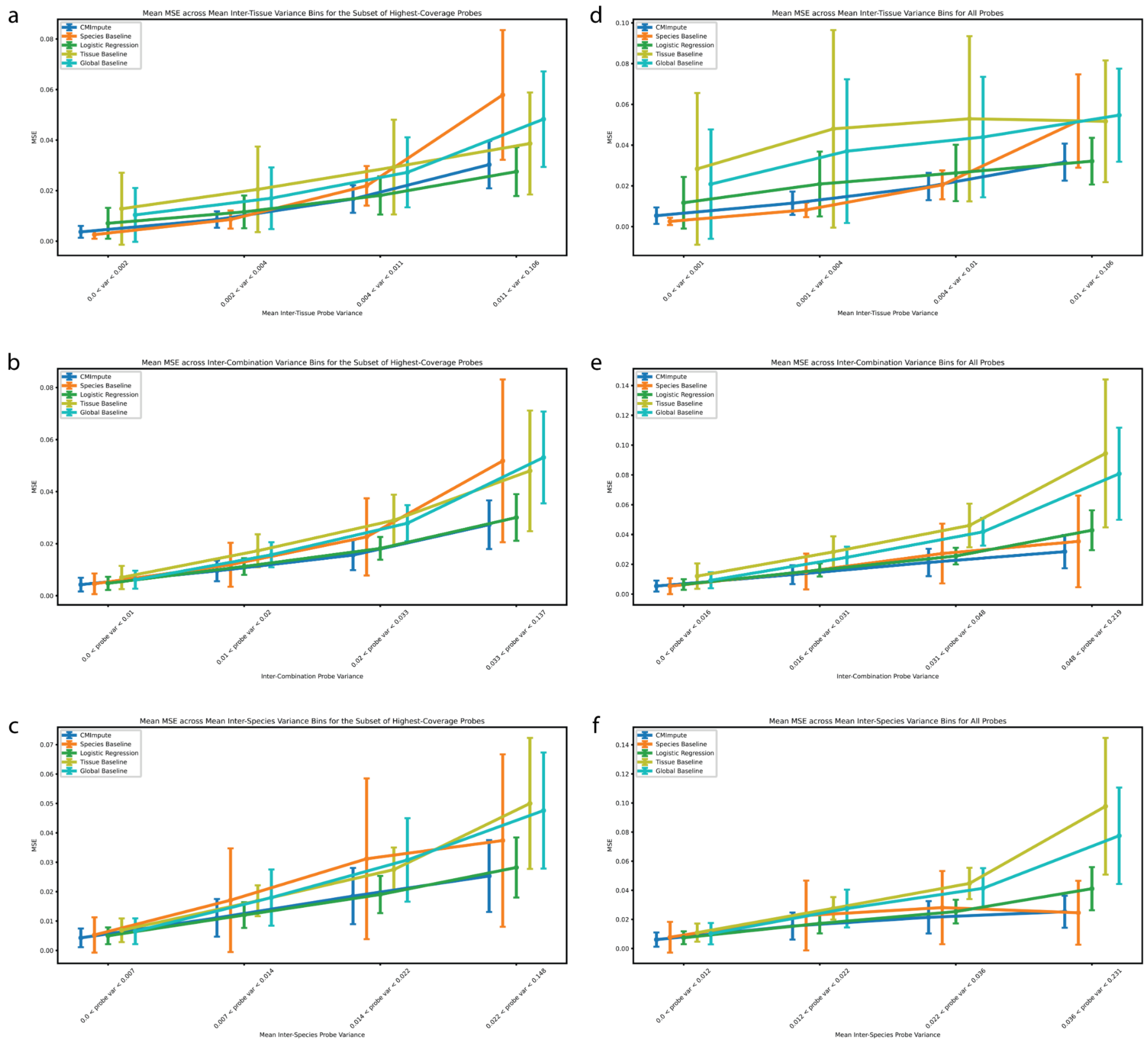


**Supplementary Figure 11. Mean probe-wise MSE across probe variance bins. a-c)** Plot of mean probe-wise MSE across **a)** mean inter-tissue, **b)** inter-combination, and **c)** mean inter-species variance quartiles when considering the subset of highest-coverage probes. Probes are divided into four bins based on ascending variance. Error bars represent one standard deviation of the probe-wise MSE within each quartile. **d-f)** Same as a-c but when considering all probes.

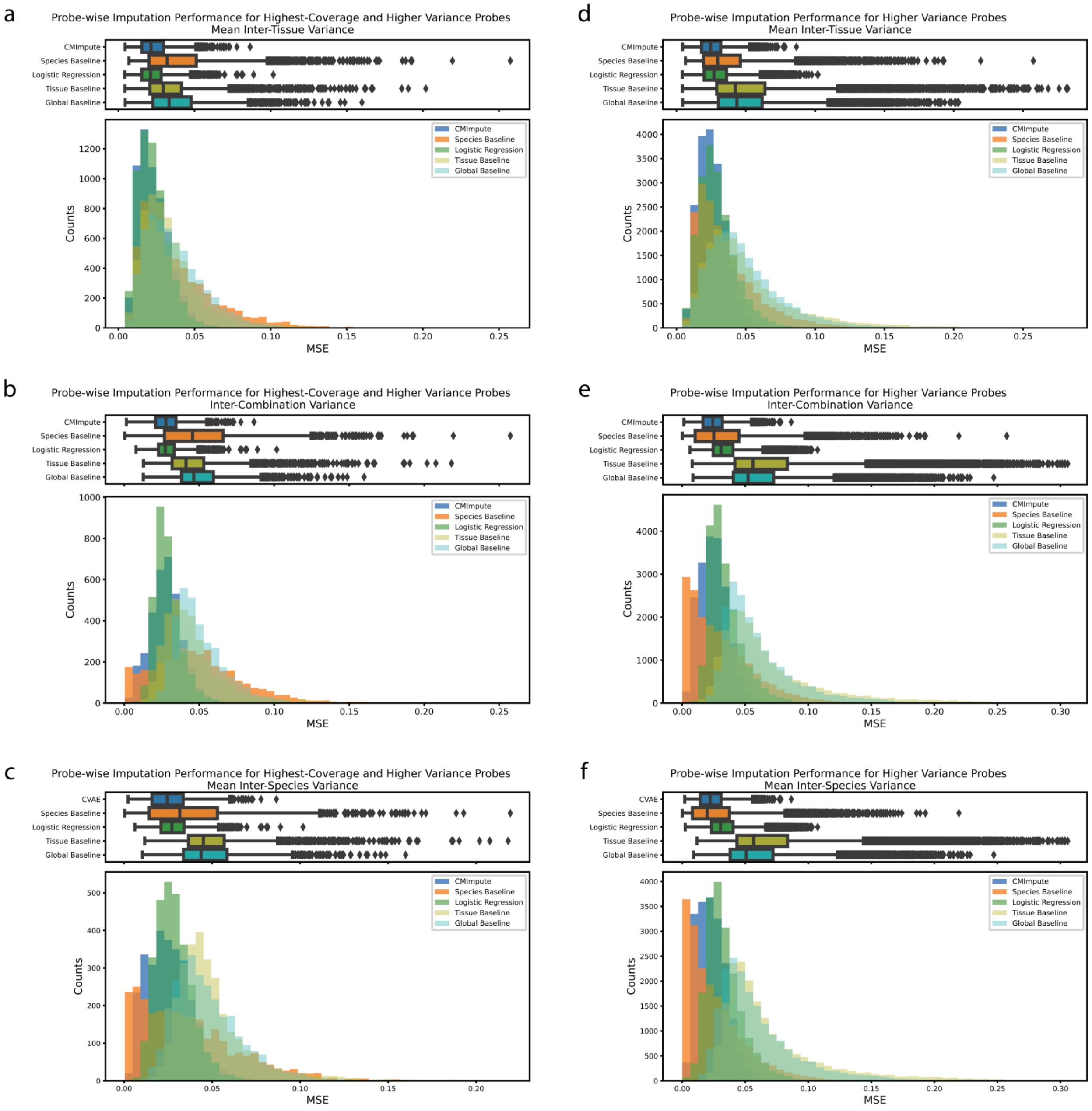


**Supplementary Figure 12. Probe-wise performance distributions for higher variance probes.** Analogous to Pearson correlation distributions in Supplementary Fig. 11. Distributions of probe-wise MSE with held-out observed values for **a)** 5,997 probes that are among the set of the highest-coverage probes and have a mean inter-tissue variance > 0.004 (median mean inter-tissue variance among set of highest-coverage probes), **b)** 3,390 probes that are among the set of the highest-coverage probes and have an inter-combination variance > 0.031 (median inter-combination variance among set of highest-coverage probes), **c)** 2,883 probes that are among the set of the highest-coverage and have a mean inter-species variance > 0.022 (median mean inter-species variance among set of highest-coverage probes), **d)** 18,746 probes with a mean inter-tissue variance > 0.004 (median mean inter-tissue variance), **e)** 18,746 probes with an inter-combination variance > 0.031 (median inter-combination variance), and **f)** 18,746 probes with a mean inter-species variance > 0.022 (median mean inter-species variance). The top boxplots show the distribution of probe-wise correlations with held-out observed values. The bottom boxplots show the number of imputed combination mean samples across 50 performance bins. Legend for both boxplots and histograms shown in histogram plot and indicate which method (CMImpute and all baselines) are being considered.

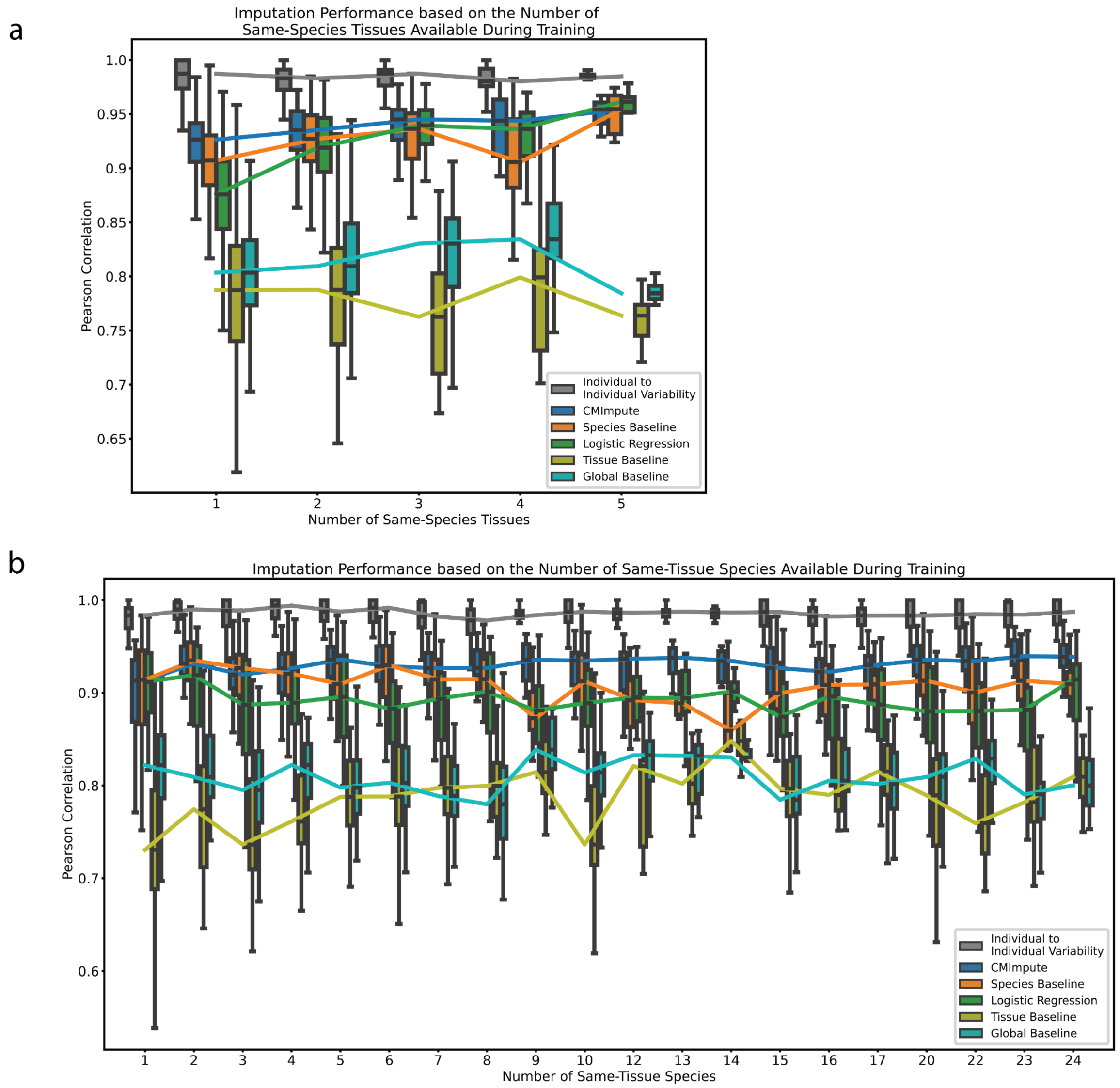


**Supplementary Figure 13. Impact of the amount of available training data on CMImpute and baseline performance. a)** Sample-wise Pearson correlation based on the number of tissue types available in the target species during training. The box plot shows the distribution of Pearson correlation for each number of tissue types. Line connects the median correlations for an imputation method across all tissue type counts. Individual to individual variability represents the average pairwise correlation of observed data between individuals of the same species and tissue type for each combination. **b)** Sample-wise Pearson correlation based on the number of species available during training in the target tissue.

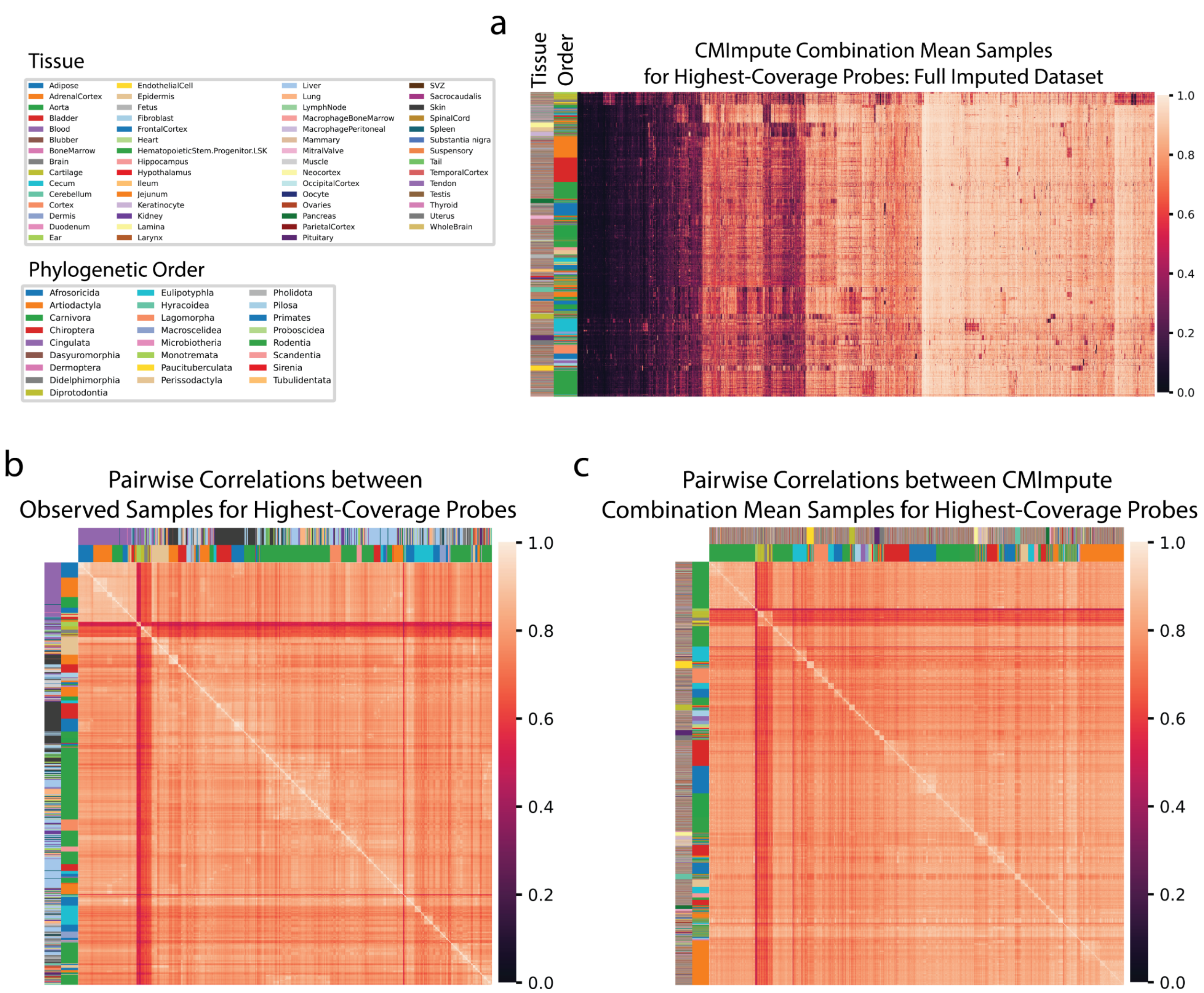


**Supplementary Figure 14. Visualization of highest-coverage probes for CMImpute's full imputed dataset. a)** Heatmap of imputed dataset's methylation probe values when considering the subset of highest-coverage probes. Each row is an imputed species-tissue combination mean sample and each column is a methylation probe restricted to the subset of highest-coverage probes. Samples and probes ordered hierarchical clustering followed by optimal leaf ordering. Samples labeled via color bar by phylogenetic order (inner) and tissue (outer). **b-c)** Similar heatmap of pairwise correlations to Fig. 6b-c for **b)** the 746 observed species-tissue combinations or **c)** 20,251 CMImpute-imputed species-tissue combinations when restricted to the subset of highest-coverage probes.

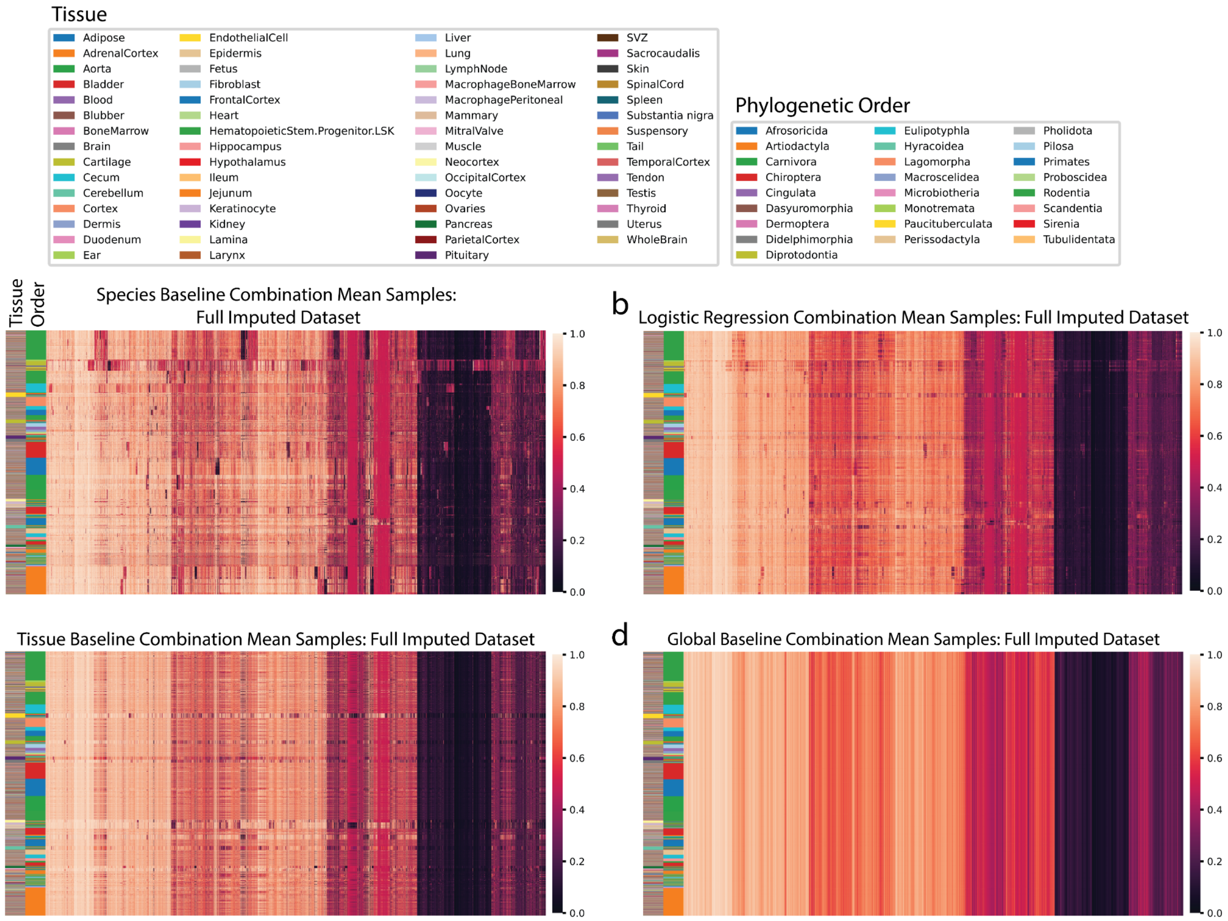


**Supplementary Figure 15. Visualization of baseline-imputed samples of non-observed combinations. a-d)** Heatmaps of **a)** species baseline, **b)** logistic regression, **c)** tissue baseline, and **d)** global baseline-imputed datasets' methylation probe values corresponding to the CMImpute-imputed dataset displayed in Fig. 6a. Samples and probes ordered hierarchical clustering followed by optimal leaf ordering. Color bars on the left indicate the phylogenetic order (inner) and tissue (outer) corresponding to the samples.

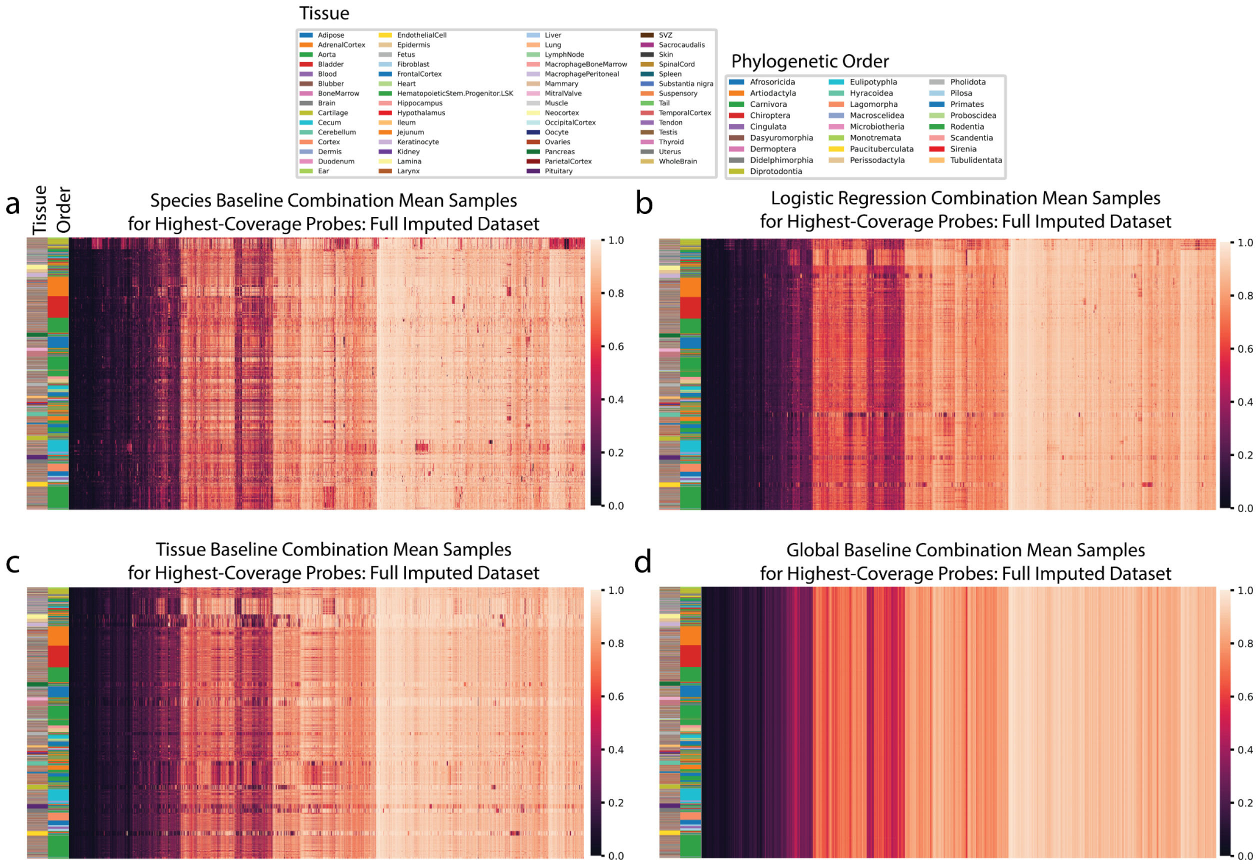


**Supplementary Figure 16. Visualization of baseline-imputed samples of non-observed combinations on the subset of highest-coverage probes. a-d)** Heatmaps of **a)** species baseline, **b)** logistic regression, **c)** tissue baseline, and **d)** global baseline-imputed datasets' methylation probe values when considering the subset of highest-coverage probes corresponding to the CMImpute-imputed dataset displayed in Supplementary Fig. 14a. Samples and probes ordered hierarchical clustering followed by optimal leaf ordering. Color bars on the left indicate the phylogenetic order (inner) and tissue (outer) corresponding to the samples.

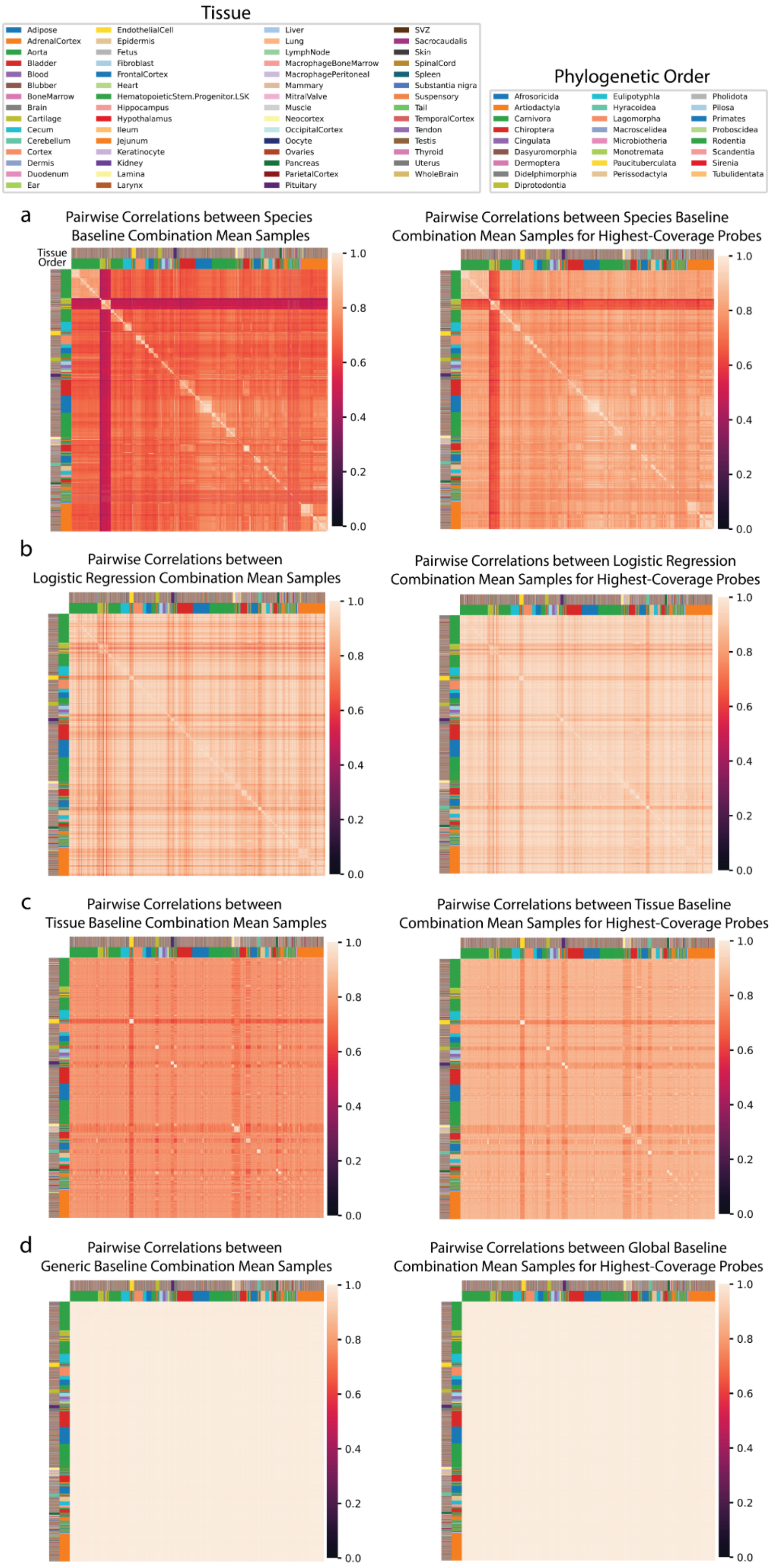
Tissue
Adipose
AdrenalCortex
Aorta
Bladder
Blood
Blubber
BoneMarrow
Brain
Cartilage
Cecum
Cerebellum
Cortex
Dermis
Duodenum
Ear
EndothelialCell
Epidermis
Fetus
Fibroblast
FrontalCortex
Heart
HematopoieticStem.Progenitor.LSK
Hippocampus
Hypothalamus
Ileum
Jejunum
Keratinocyte
Kidney
Lamina
Larynx
Liver
Lung
LymphNode
MacrophageBoneMarrow
MacrophagePeritoneal
Mammary
MitralValve
Muscle
Neocortex
OccipitalCortex
Oocyte
Ovaries
Pancreas
ParietalCortex
Pituitary
SVZ
Sacrocaudalis
Skin
SpinalCord
Spleen
Substantia nigra
Suspensory
Tail
TemporalCortex
Tendon
Testis
Thyroid
Uterus
WholeBrain
Phylogenetic Order
Afrosoricida
Artiodactyla
Carnivora
Chiroptera
Cingulata
Dasyuromorphia
Dermoptera
Didelphimorphia
Diprotodontia
Eulipotyphla
Hyracoidea
Lagomorpha
Macroscelidea
Microbiotheria
Monotremata
Paucituberculata
Perissodactyla
Pholidota
Pilosa
Primates
Proboscidea
Rodentia
Scandentia
Sirenia
Tubulidentata
a
Pairwise Correlations between Species Baseline Combination Mean Samples
Tissue
Order
Pairwise Correlations between Species Baseline Combination Mean Samples for Highest-Coverage Probes
b
Pairwise Correlations between Logistic Regression Combination Mean Samples
Pairwise Correlations between Logistic Regression Combination Mean Samples for Highest-Coverage Probes
c
Pairwise Correlations between Tissue Baseline Combination Mean Samples
Pairwise Correlations between Tissue Baseline Combination Mean Samples for Highest-Coverage Probes
d
Pairwise Correlations between Generic Baseline Combination Mean Samples
Pairwise Correlations between Global Baseline Combination Mean Samples for Highest-Coverage Probes
1.0
0.8
0.6
0.4
0.2
0.0

**Supplementary Figure 17. Visualization of species and tissue signals via pairwise correlations for baselines. a-d)** Similar heatmaps as in Fig. 6c and Supplementary Fig. 14c here showing pairwise correlations between 20,251 **a)** species baseline, **b)** logistic regression, **c)** tissue baseline, and **d)** global baseline-imputed combination mean samples when considering all probes (left) and the subset of highest-coverage probes (right). Samples are ordered based on hierarchical clustering followed by optimal leaf ordering of the full methylation samples from each dataset. Color bars on the left indicate the phylogenetic order (inner) and tissue (outer) corresponding to the samples.

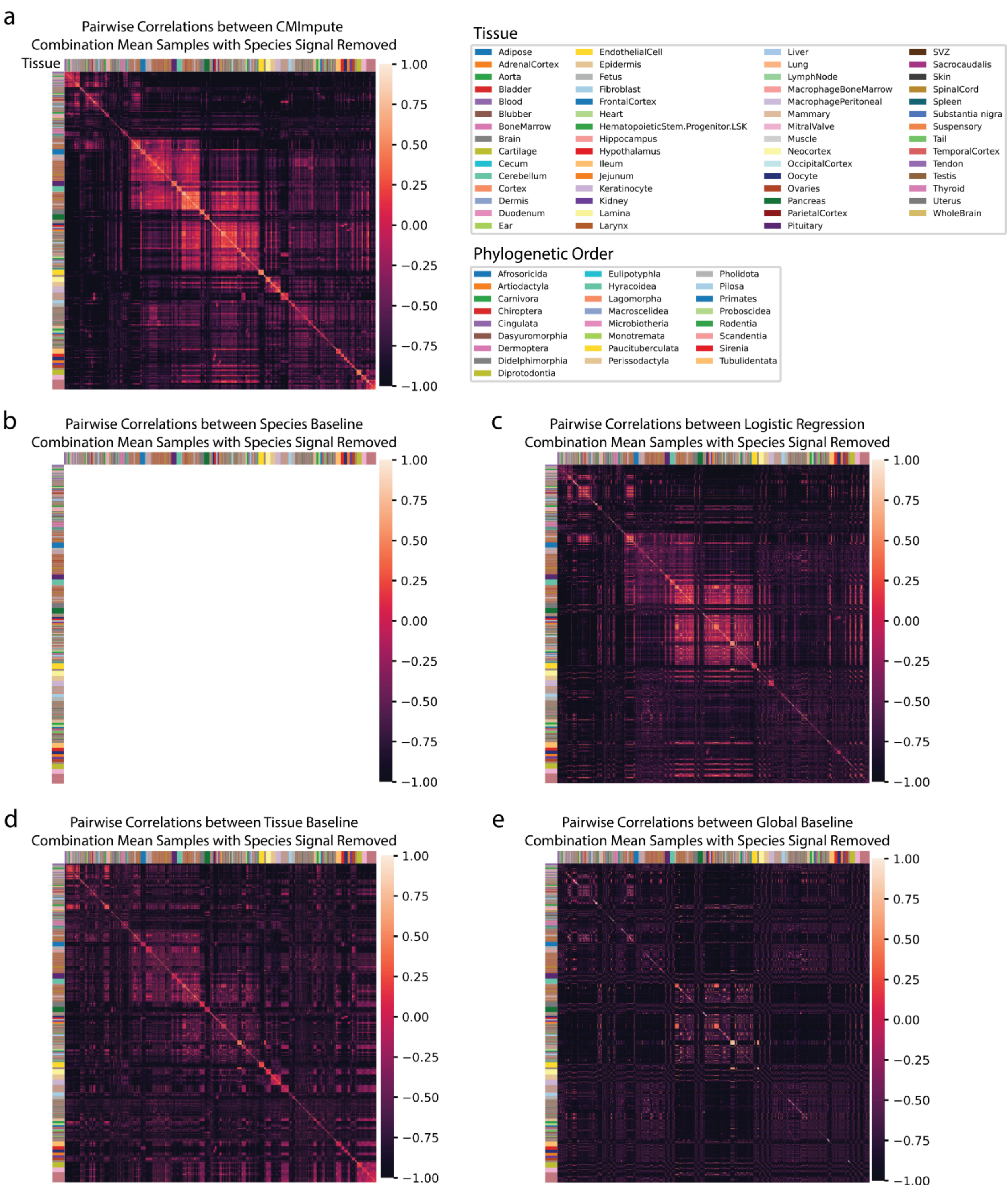


**Supplementary Figure 18. Visualization of tissue signal in imputed samples of non-observed combinations. a-e)** Heatmaps of pairwise correlations between 20,251 **a)** CMImpute, **b)** species baseline, **c)** logistic regression, **d)** tissue baseline, and **e)** global baseline-imputed combination mean samples with the species signal removed (as shown in Fig. 6d). Samples are ordered based on hierarchical clustering of the difference between the full imputed samples and the species baseline samples followed by optimal leaf ordering.

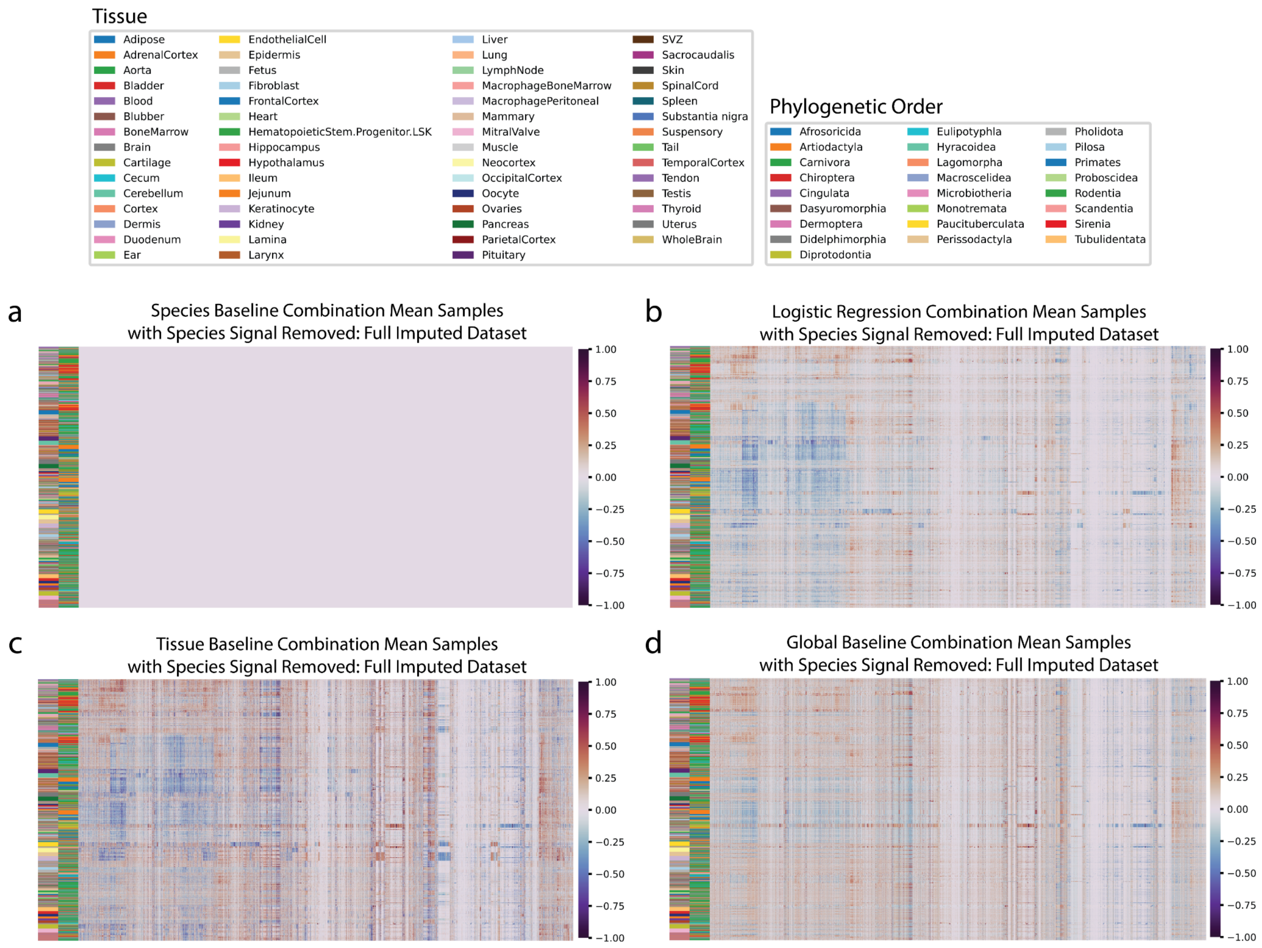


**Supplementary Figure 19. Visualization of baseline-imputed combination mean samples with the species signal removed. a-d)** Heatmaps of **a)** species baseline, **b)** logistic regression, **c)** tissue baseline, and **d)** global baseline-imputed combination mean samples with the species signal removed. Sample and probe order, color bar labels, and color scale corresponds to Fig. 6d.

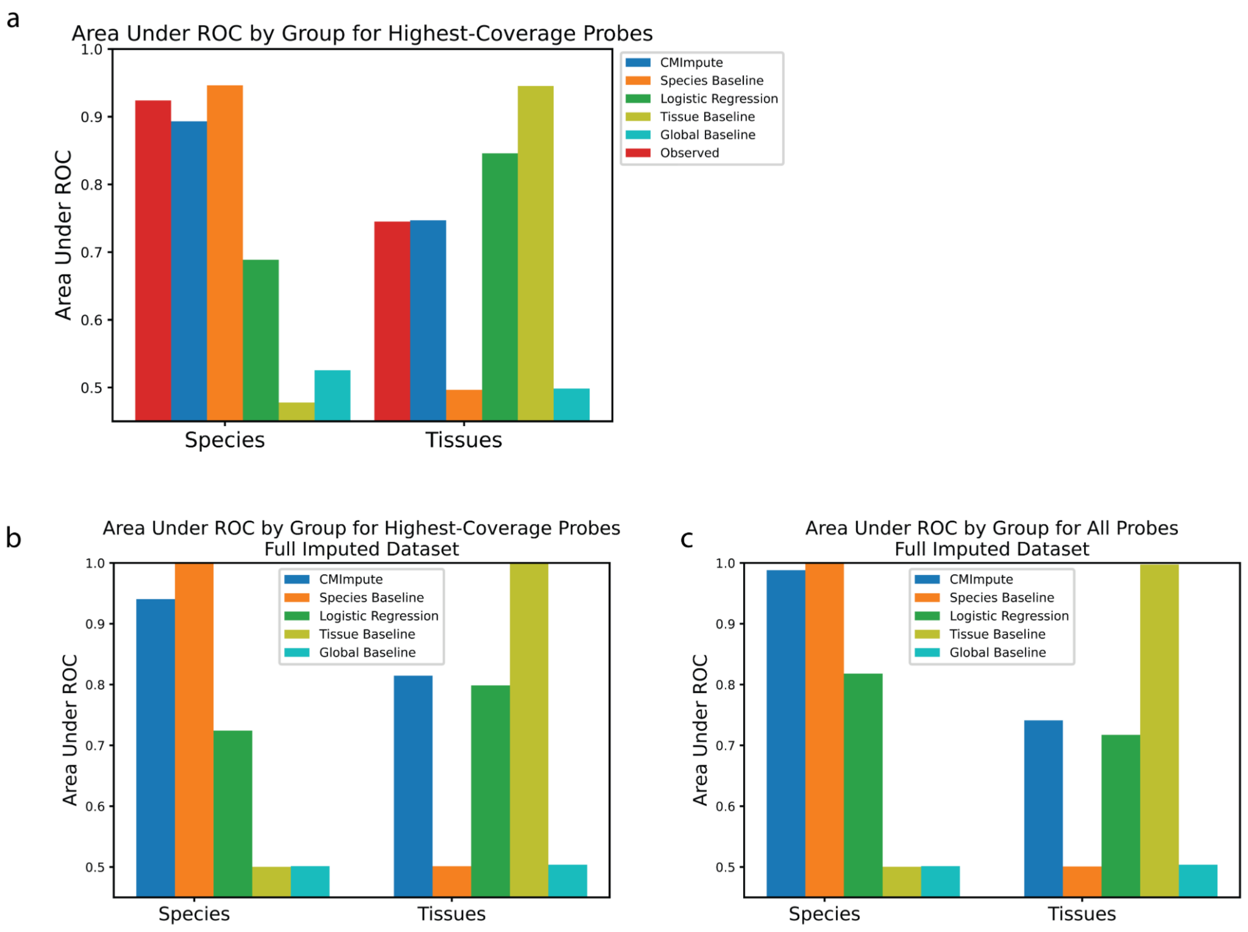


**Supplementary Figure 20. Species and tissue signal for the subset of highest-coverage probes and the full imputed dataset. a)** Area Under ROC values for predicting whether samples within the cross-validation dataset are from the same species (left) or tissue (right) based on their pairwise correlations for the subset of highest-coverage probes. Results when considering all probes shown in Fig. 7. **b-c)** Area Under ROC values for predicting whether samples within the full 20,251 combination mean sample imputed dataset are from the same species (left) or tissue (right) based on their pairwise correlations when considering **b)** the subset of highest-coverage probes and **c)** all probes.

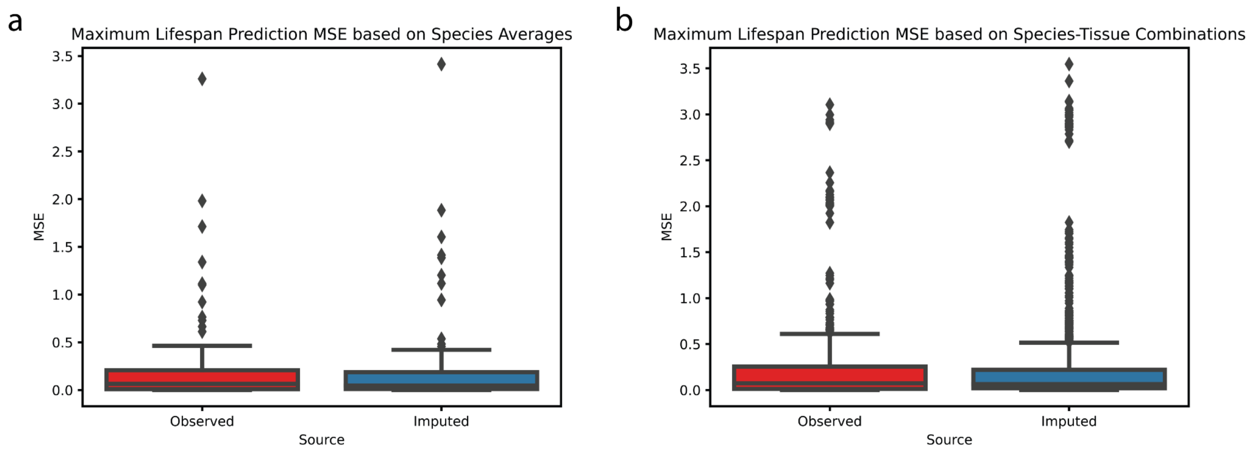


**Supplementary Figure 21. Prediction of species' maximum lifespan using combination mean samples from different tissue types. a)** Boxpot comparing the log-maximum lifespan prediction MSE using observed vs. imputed averages of combination mean samples within each species. Both the imputed and experimentally-profiled combination mean samples span the same 114 species. **b)** Same as a but comparing the log-maximum lifespan prediction mean squared error using observed vs. imputed species-tissue combination mean samples. The 441 observed combination mean samples span all observed tissues and the 6,285 imputed combination mean samples span all non-observed tissues across the 114 species.

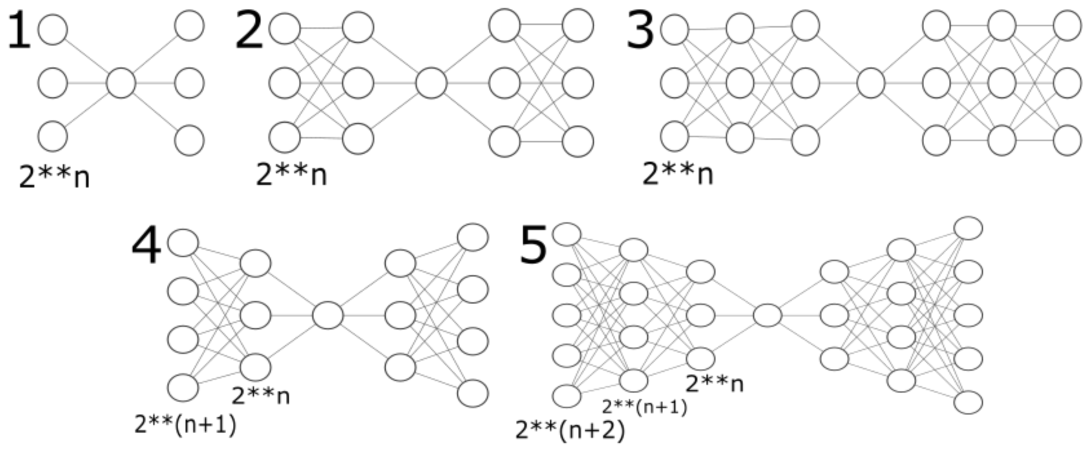


**Supplementary Figure 22. Potential network architectures.** Five network architectures that CMImpute considered during the hyperparameter grid search. Each architecture is symmetrical around the latent space with equal numbers of hidden layers in the encoder and decoder. The layout options consist of one through three hidden layers in the encoder and decoder and both tapered (each layer closer to the latent space gets smaller) and equal dimension (each layer in the encoder and decoder is the same dimension) for two and three hidden layer options. Each option (1-5) corresponds to the layouts parameter and n corresponds to the hidden layer dimension parameter (n) from Supplementary Table 2. The final network layout, along with all other hyperparameters, are selected via hyperparameter grid search. Notation of 2**n represents 2 to the power of n.

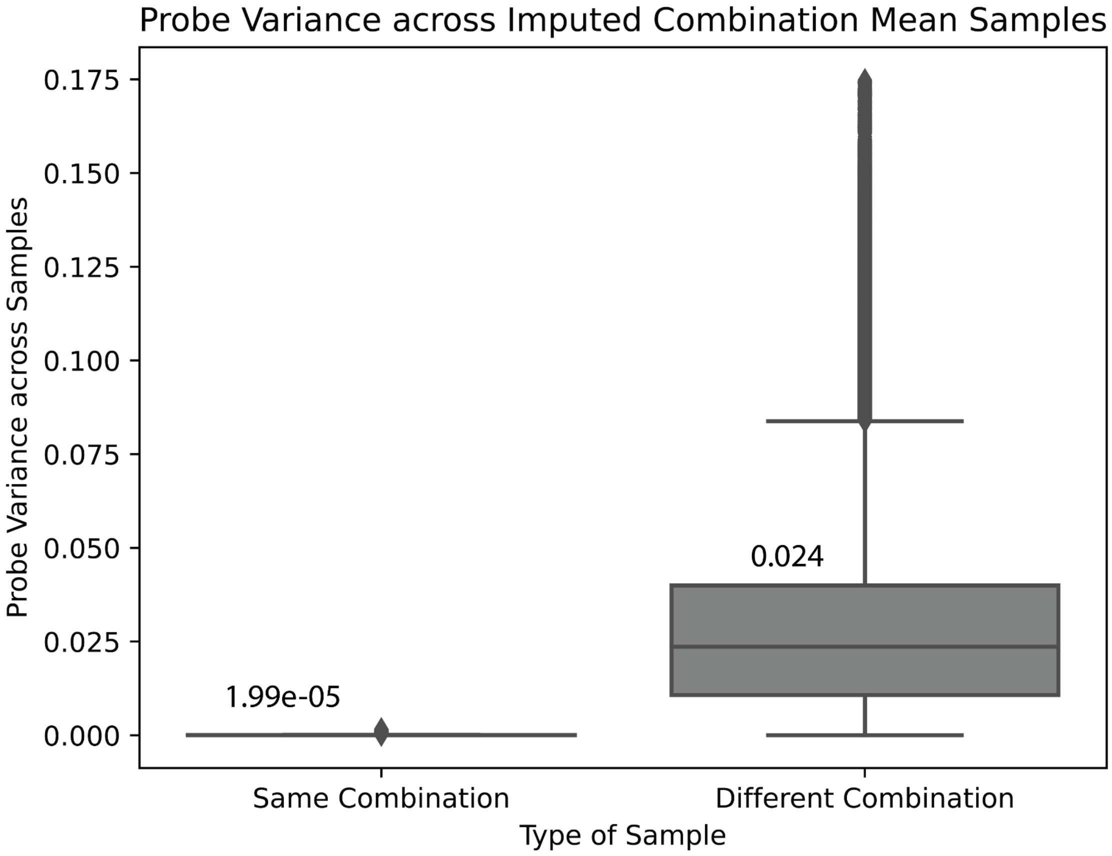


**Supplementary Figure 23. Impact of the random normal latent space sampling on the final imputation result.** For a random fold, each held-out combination is imputed 20 times using a different random normal sampling for the latent representation. For each species-tissue combination, the variance across the 20 samples with different random normal latent samplings was calculated. Additionally, the variance across samples from different species-tissue combinations was calculated. The boxplot shows the difference in variance between samples of the same combination but different latent samplings and samples of different combinations. Each box labeled with median probe variance.

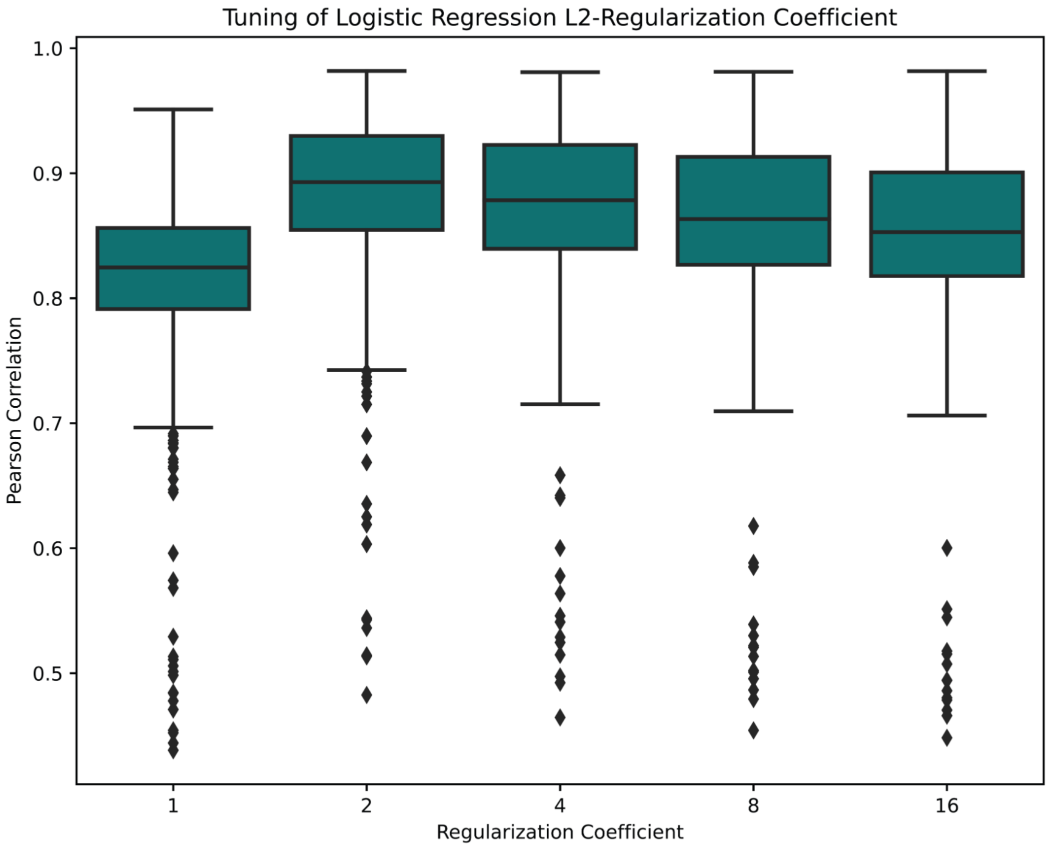


**Supplementary Figure 24. Tuning of the logistic regression baseline $L_2$-regularization coefficient.** For each coefficient, a logistic regression model was trained for each probe on each of the five cross-validation folds (Methods). The boxplot shows the validation sample-wise Pearson correlation with held-out observed values for regression coefficients of 1, 2, 4, 8, and 16.

| | Pearsonr All Probes | MSE All Probes | Pearson r Highest-Coverage Probes | MSE Highest-Coverage Probes |
|---|---|---|---|---|
| Species Baseline | 68% | 68% | 83% | 82% |
| Logistic Regression | 78% | 74% | 65% | 59% |
| Tissue Baseline | 98% | 98% | 93% | 92% |
| Global Baseline | 97% | 97% | 94% | 92% |

**Supplementary Table 1. Percentage of imputed samples where CMImpute outperforms baselines.** For each baseline method (species baseline, logistic regression, tissue baseline, and global baseline top to bottom), the table displays the percentage of imputed species-tissue combination mean samples from the cross-validation analysis where CMImpute outperforms the baseline. A percentage is reported based on Pearson correlation considering all probes, MSE considering all probes, Pearson correlation restricted to the subset of highest-coverage probes, and MSE restricted to the subset of highest-coverage probes (left to right).

| Hyperparameter | Option |
| --- | --- |
| n | 8, 9, 10, 11 |
| Layout | 1, 2, 3, 4, 5 |
| Activation Function | ReLu, Sigmoid, TanH |
| Learning Rate | 0.001, 0.01 |
| Epsilon | 1e-7, 1e-5, 0.001, 0.1 |
| Latent Space Dimension | 2, 4, 8 |

**Supplementary Table 2. Hyperparameters for the CVAE model.** Search space for the grid search consists of all potential combinations of hyperparameter values. n is the parameter used to calculate the hidden layer dimensions and Layout corresponds to the network architecture in Supplementary Fig. 22. Activation function used to initialize each hidden layer. Learning Rate and Epsilon are parameters in the Adam optimizer. Latent space dimension is the dimension of the latent layer.
